# DNMT3A haploinsufficiency results in behavioral deficits and global epigenomic dysregulation shared across neurodevelopmental disorders

**DOI:** 10.1101/2020.07.10.195859

**Authors:** Diana L. Christian, Dennis Y. Wu, Jenna R. Martin, J. Russell Moore, Yiran R. Liu, Adam W. Clemens, Sabin A. Nettles, Nicole M. Kirkland, Cheryl A. Hill, David F. Wozniak, Joseph D. Dougherty, Harrison W. Gabel

## Abstract

Mutations in DNA methyltransferase 3A (DNMT3A) have been detected in autism and related disorders, but how these mutations disrupt nervous system function is unknown. Here we define the effects of neurodevelopmental disease-associated DNMT3A mutations. We show that diverse mutations affect different aspects of protein activity yet lead to shared deficiencies in neuronal DNA methylation. Heterozygous DNMT3A knockout mice mimicking DNMT3A disruption in disease display growth and behavioral alterations consistent with human phenotypes. Strikingly, in these mice we detect global disruption of neuron-enriched non-CG DNA methylation, a binding site for the Rett syndrome protein MeCP2. Loss of this methylation leads to enhancer and gene dysregulation that overlaps with models of Rett syndrome and autism. These findings define effects of DNMT3A haploinsufficiency in the brain and uncover disruption of the non-CG methylation pathway as a convergence point across neurodevelopmental disorders.

## Introduction

Precise regulation of transcription through epigenetic mechanisms is critical for nervous system development (Cholewa-Waclaw et al., 2016). Exome sequencing studies have revealed mutations of genes encoding epigenetic modifiers of chromatin structure as a major underlying cause of neurodevelopmental diseases (NDD), including autism spectrum disorder (ASD) (McRae et al., 2017; Sanders et al., 2015; Satterstrom et al., 2019). A challenge emerging from these discoveries is to define the cellular functions of the disrupted proteins during normal development and search for shared pathways between these proteins that can potentially be targeted for therapeutic development.

Gene regulation mediated by DNA methylation has emerged as an epigenetic mechanism that plays a critical role in nervous system function (Kinde et al., 2015). In addition to the classical methylation of cytosines found at CG dinucleotides (mCG), neurons contain uniquely high levels of methyl-cytosine (mC) in a non-CG context, with this mark occurring primarily at CA dinucleotides (mCA) (Guo et al., 2014; Lister et al., 2013a; Xie et al., 2012). mCA is deposited *de novo* through the activation of the DNA methyltransferase 3A (DNMT3A) enzyme during the early postnatal period (1-6 weeks of age in mice). Levels of mCA increase specifically in neurons until the number of methylation sites in the non-CG context are nearly equivalent to mCG sites (Guo et al., 2014; Lister et al., 2013a; Xie et al., 2012). A critical function for mCA is to serve as a binding site for a neuron-enriched chromatin protein, Methyl-CpG binding Protein 2 (MeCP2) (Chen et al., 2015; Gabel et al., 2015; Guo et al., 2014). MeCP2 was initially defined by its high affinity for mCG, but biochemical and genomic studies indicate that it preferentially interacts with mCA to down-regulate transcription of genes with essential functions in the brain (Boxer et al., 2019; Gabel et al., 2015; Kinde et al., 2016; Lagger et al., 2017; Lyst and Bird, 2015). Loss of MeCP2 leads to the severe neurological disorder Rett syndrome, while duplication causes MeCP2-duplication syndrome, an ASD, suggesting that read-out of mCA is critical to nervous system function (Amir et al., 1999; Van Esch et al., 2005).

Notably, human exome sequencing studies have recently identified *de novo* mutations in DNMT3A in individuals with ASD (Feliciano et al., 2019; Sanders et al., 2015; Satterstrom et al., 2019). Separate studies have also defined heterozygous disruption of DNMT3A as the underlying cause of Tatton-Brown Rahman syndrome (TBRS), a heterogeneous NDD characterized by intellectual disability, overgrowth, craniofacial abnormalities, anxiety, and high penetrance of ASD (Tatton-Brown et al., 2014, 2018). While a portion of the mutations identified in affected individuals are truncations that are predicted to cause complete inactivation of the enzyme, a majority of disease-associated alleles are missense mutations, raising questions about whether loss-of-function effects are a primary mechanism of disruption in DNMT3A disorders (Tatton-Brown et al., 2014, 2018). In addition, while heterozygous loss of DNMT3A has been studied in the context of oncogenesis in the hematopoietic system (Cole et al., 2017), the effects of partial loss of DNMT3A on nervous system function *in vivo* have not been examined; therefore, the consequences of possible methylation changes on neuronal gene regulation and behavior are unknown.

Here we examine the molecular effects of neurodevelopmental disease-associated DNMT3A mutations and explore the consequences of heterozygous DNMT3A mutation on the neuronal epigenome. Our results indicate that missense mutations across canonical domains of DNMT3A disrupt different aspects of protein function, yet mutations in all domains reduce the capacity of the enzyme to deposit neuronal mCA. We detect altered growth and behavior in DNMT3A heterozygous deletion mice, supporting haploinsufficiency as a driver of pathology in DNMT3A disorders. Through integrated epigenomic analysis, we reveal disruption of mCA throughout the brain of DNMT3A mutant mice. Strikingly, we show that this loss of mCA leads to disruption of distal regulatory enhancer activity and changes in gene expression that overlap with models of MeCP2 disorders and other ASDs. These findings define the effects of NDD-associated DNMT3A mutations for the first time and reveal disruption of mCA-mediated epigenomic regulation as a convergence site across clinically distinct NDDs.

## Results

### Functional analysis of disease-associated DNMT3A mutations

Multiple DNMT3A mutations have been identified in individuals with ASD and TBRS. However, the large number of missense mutations identified and the phenotypic heterogeneity of individuals with these mutations raise the possibility that alterations of amino acids within different protein domains may have distinct consequences and may dictate the nature and severity of disease. We therefore sought to assess the effects of disease-associated DNMT3A mutations on protein expression, cellular localization, and catalytic activity, looking for common effects that may be core to the development of NDD.

We engineered amino-acid alterations homologous to human disease mutations into a FLAG-tagged DNMT3A protein expression vector and assessed multiple mutations found within each functional domain of the protein (Figure 1A). These analyses included mutations in the chromatin interacting proline-tryptophan-tryptophan-proline (PWWP) domain, the auto-inhibitory Histone H3 lysine 4 interacting ATRX-DNMT3-DNMT3L (ADD) domain, and the well-defined methyltransferase catalytic domain (Gowher and Jeltsch, 2018). Transfection into heterologous cells facilitated rapid assessment of protein expression by western blot, cellular localization by immunocytochemistry, and catalytic activity using an *in vitro* methyltransferase assay (Figure 1). Mutations in the PWWP domain resulted in a reduction in DNMT3A protein expression and loss of nuclear localization compared to wild-type controls (Figure 1B-D, Figure S1A-C). When expressed at equal levels to that of wild-type protein however, these mutations exhibited substantial catalytic activity (Figure 1E,F). In contrast, mutations found in the catalytic methyltransferase domain of DNMT3A showed wild-type expression and localization but displayed deficits in catalytic activity in the *in vitro* methyltransferase analysis (Figure 1B-F, Figure S1A-C). Mutations in the ADD domain of DNMT3A displayed normal protein localization and expression levels and exhibited equal or higher methylation activity compared to wild-type protein *in vitro* (Figure 1B-F, Figure S1A-C).

**Figure 1.**
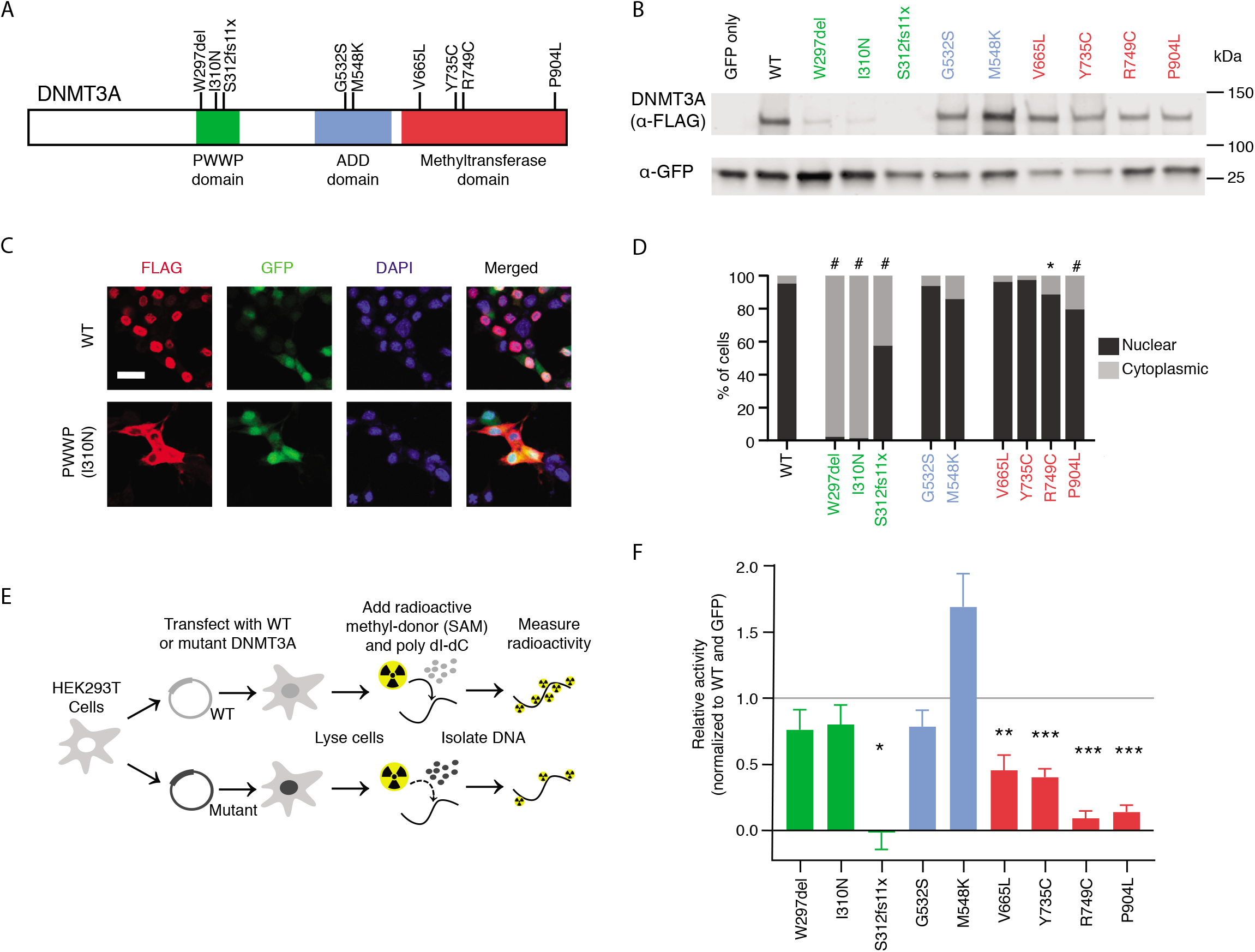
Disease-associated DNMT3A mutations disrupt distinct aspects of protein function. (**A**) Schematic of human DNMT3A protein showing canonical domains and disease-associated mutations identified in previous studies (Sanders et al., 2015; Tatton-Brown et al., 2018). (**B**) Example immunoblot of DNMT3A mutant protein expression. (**C**) Example images of DNMT3A protein immunocytochemistry from wild type and PWWP domain mutant. Scale bar = 20μm. (**D**) Quantification of DNMT3A mutant protein localization (#, *P*<0.0001; *, *P*<0.05; n=6-16 images; Generalized Linear Model test of percent nuclear expression per image for mutants compared to WT with Bonferroni correction). (**E**) Schematic of *in vitro* methylation assay for DNMT3A mutant proteins. (**F**) Activity of DNMT3A mutant proteins in the *in vitro* methylation assay. (***, *P*<0.001; **, *P*<0.01; *, *P*<0.05; n=4-19; one-sample Student’s T-Test from normalized WT mean of 1 with Bonferroni correction). Bar graphs indicate mean with SEM error bars.

To further evaluate the functional effects of disease-associated DNMT3A mutations in the context of endogenous chromatin, we tested the capacity of DNMT3A mutants to establish DNA methylation across the genome in mouse cortical neurons. For this analysis, we focused on the global build-up of mCA in postmitotic neurons that requires DNMT3A (Gabel et al., 2015; Lister et al., 2013a). Cultured neurons isolated from the cerebral cortex at embryonic day 14.5 accumulate mCA *in vitro* and this build-up can be blocked by lentiviral-mediated delivery of Cre recombinase to DNMT3A^flx/flx^ cells at 3 days *in vitro* (DIV) (Figure 2A, Figure S1D,E). We cotransduced wild-type or mutant DNMT3A lentivirus at equal levels (Figure S1F) to test the capacity of each protein to rescue deposition of DNA methylation. Analysis of mutations across the major domains of DNMT3A detected deficits in mCA accumulation for all disease-associated mutations tested (Figure 2B). Notably, mutations in the ADD domain that exhibited robust catalytic activity *in vitro* displayed moderate-to-severe deficits in mCA deposition in neurons. The ADD domain has been implicated in both histone binding and auto-inhibition of the protein (Guo et al., 2015), thus effects in this neuronal assay may indicate that loss of ADD function blocks the capacity of the enzyme to engage with chromatin and promote DNMT3A methylation activity in cells. Together our results indicate that although NDD-associated mutations in DNMT3A affect different protein domains and alter distinct aspects of protein function (e.g. localization, chromatin interaction, catalysis), these mutations share a common outcome of reduced enzymatic activity on neuronal DNA, with many mutations resulting in functionally null proteins.

**Figure 2.**
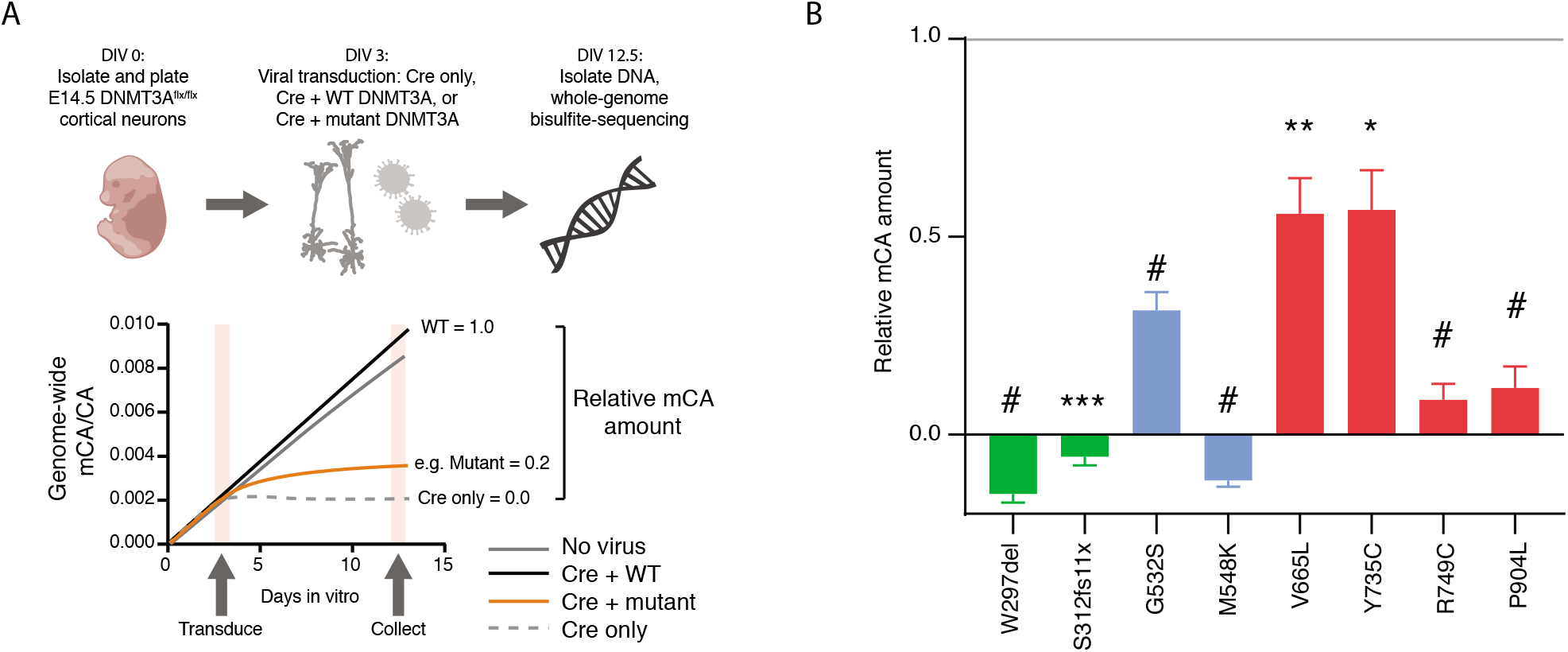
Disease-associated DNMT3A mutations prevent buildup of neuronal CA methylation. (**A**) Schematic of DNMT3A functional analysis in primary culture neurons. Cortical neurons are collected from DNMT3A^flx/flx^ mice at E14.5 and cultured. After 3 days *in vitro* (DIV), neurons are virally transduced with Cre recombinase and WT or mutant FLAG-tagged DNMT3A. On DIV 12.5, DNA and RNA are collected. Equal DNMT3A mRNA expression is verified by qRT-PCR (Figure S1D) and DNA is used for whole genome bisulfite sequencing analysis. (**B**) Relative mCA amount compared to Cre only and Cre+WT DNMT3A controls (#, *P*<0.0001; ***, *P*<0.001; **, *P*<0.01; *, *P*<0.05; n=4-11; one-sample Student’s T-Test from normalized WT mean of 1 with Bonferroni correction). Bar graphs indicate mean with SEM error bars.

### *In vivo* effects of heterozygous DNMT3A disruption

In light of our findings that multiple NDD-associated missense mutations in DNMT3A result in complete or near-complete loss of function, we next sought to understand the effects of heterozygous inactivation of DNMT3A *in vivo*. Previous studies have demonstrated severe developmental deficits and perinatal lethality associated with complete loss of DNMT3A (homozygous null mutation) in mice (Okano et al., 1999). However, the relevance of heterozygous mutation of DNMT3A to neurodevelopmental disease has only recently been uncovered and growth and behavioral effects of partial DNMT3A inactivation have not been systematically assessed. We therefore carried out growth, behavioral, and molecular analyses of mice carrying a constitutive heterozygous deletion of exon 19 of *Dnmt3a* (see *methods*) (Kaneda et al., 2004). We find that this mutation leads to 50% reduction of RNA and protein expression, allowing us to study the *in vivo* effects of heterozygous null mutation of DNMT3A (referred to as DNMT3A^KO/+^) (Figure S2A-C).

We first examined phenotypes with relevance to the overgrowth in individuals with heterozygous DNMT3A mutations (Tatton-Brown et al., 2018), including enlarged body size and obesity (body weight), tall stature (long-bone length), and macrocephaly (skull dimensions). DNMT3A^KO/+^ mice showed similar body weight to controls in the early postnatal period but were significantly heavier than controls as mature adults (Figure 3A). This phenotype mimics a maturity-associated trend toward increasing body weight observed in TBRS patients (Tatton-Brown et al., 2018). Measurements of bone length indicated a small but significant increase in tibia length in DNMT3A^KO/+^ mice, with a trend towards longer femur length (Figure 3B, Figure S3A-C). Morphometric analyses of the cranium and mandible indicated very subtle differences between DNMT3A^KO/+^ mice and their wild-type littermates (Figure S3D-F). One linear distance spanning the rostrocaudal length of the interparietal bone is larger in DNMT3A^KO/+^ mice relative to wild-type littermates. Two linear distances in the facial region were significantly larger in wild-type mice, while all other comparisons were not significantly different (Figure S3D). This suggests very slight disruptions in growth of the facial region in DNMT3A^KO/+^ mice. Together these findings uncover effects on long bone length that mirror aspects of the human disorder, while skull development in DNMT3A^KO/+^ mice shows more limited effects. Additionally, enlarged body mass in these mice appears to mimic overgrowth and obesity detected in individuals with TBRS (Tatton-Brown et al., 2014).

**Figure 3.**
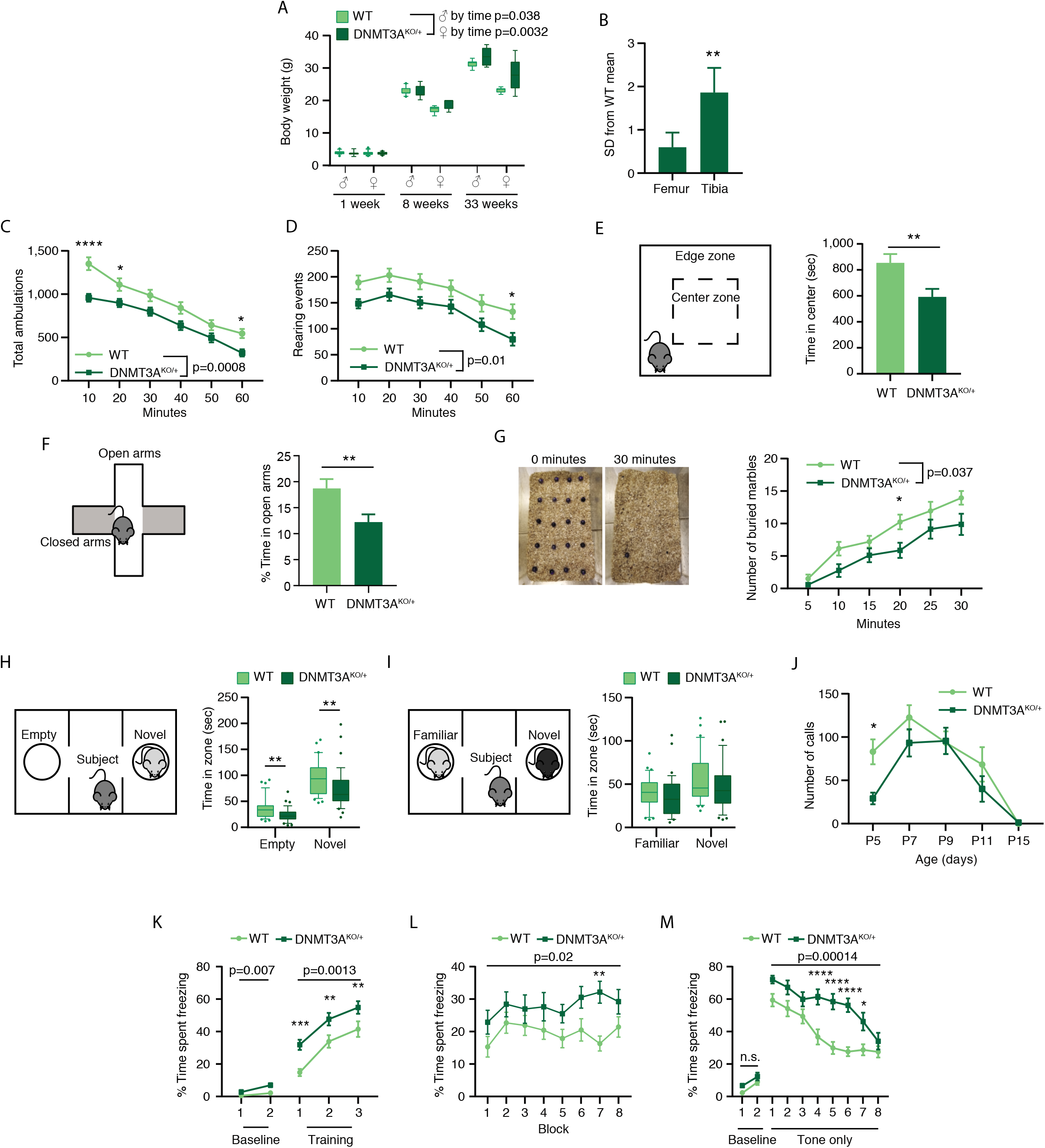
Heterozygous disruption of DNMT3A *in vivo* leads to growth and behavioral alterations. (**A**) Body weight of DNMT3A^KO/+^ and WT mice at three developmental timepoints (Male *P*=0.038 genotype by age interaction effect, F_(2,50)_=3.494, n=6-18; Female *P*=0.0032 genotype by age interaction effect, F_(2,48)_=6.498; Female *P*=0.0016 genotype effect, F_(1,48)_=11.18, n=5-17; two-way ANOVA). (**B**) Lengths of femur and tibia bones measured by dual X-ray imaging shown as standard deviations from the WT mean for the DNMT3A^KO/+^ mice (**, *P*<0.01; n=12; paired Student’s T-Test). (**C**) Total ambulations of mice during 1-hour open-field testing, split into 10-minute bins (*P*=0.0008 effect by genotype, F_(1,46)_=13.02, n=21,27; two-way repeated-measures ANOVA with Sidak’s multiple comparison test; *, *P*<0.05; ****, *P*<0.0001). (**D**) Number of rearing events of mice during 1-hour open-field testing, split into 10-minute bins (*P*=0.0103 effect by genotype, F_(1,46)_=7.161, n=21,27; two-way repeated-measures ANOVA with Sidak’s multiple comparison test; *, *P*<0.05). (**E**) Schematic of open-field testing center and edge zones (left). Total time spent in the center zone of field during open-field testing (right) (*P*=0.0075, n=21,27; unpaired Student’s T-Test). (**F**) Schematic of the elevated plus maze indicating closed and open arms (left), and percent of time mice spent in the open arms compared to all arms during first day of testing (right) (*P=*0.0069; n=33,39; unpaired Student’s T-Test). (**G**) Example images of marble burying assay (left) and quantification of marbles buried during 30 minutes of testing split into 5-minute bins (right)(*P*=0.0374 effect by genotype, F_(1,25)_=4.834, n=14,13; two-way repeated-measures ANOVA with Sidak’s multiple comparison test; *, *P*<0.05). (**H**) Schematic of 3-chamber task in which subject mouse can freely explore apparatus containing a novel mouse or an empty cup (left). Quantification of time spent in zones closest to each cup (right) (Empty, *P*=0.0026; Novel, *P*=0.0095; n=33,39; unpaired Student’s T-Test). (**I**) Schematic of 3-chamber task in which subject mouse can freely explore apparatus and interact with novel mouse or familiar mouse (left). Time spent in zones closest to novel mouse and familiar mouse (right) (Familiar, *P*=0.29; Novel, *P*=0.24; n=33,39; unpaired Student’s T-Test). (**J**) Number of ultrasonic calls from pup isolated from the nest for 3-minute testing over developmental time points (Analysis run on days 5-9 as these were timepoints in which all animals tested had data; *P*=0.0378 effect by genotype, F_(1,285)_=4.355, n=9-46; two-way ANOVA with Sidak’s multiple comparisons test; *, *P*<0.05). (**K-M**) Percent time spent freezing in (**K**) Conditioned fear training (Baseline: *P*=0.0071 effect by genotype, F_(1,50)_=7.897; Cue: *P*=0.0013 effect by genotype, F_(1,50)_=11.7; n=26; two-way repeated-measures ANOVA with Sidak’s multiple comparisons test; **, *P*<0.01; ***, *P*<0.001), (**L**) contextual fear trials (*P*=0.0215 effect by genotype, F_(1,50)_=5.633, n=26; two-way repeated-measures ANOVA), and (**M**) cued fear trials (Baseline: *P*=0.0606 effect by genotype, F_(1,50)_=3.685; Cue: *P*<0.0001 effect by genotype, F_(1,50)_=17.03; n=26; two-way repeated-measures ANOVA with Sidak’s multiple comparisons test; *, *P*<0.05; ****, *P*<0.0001). Line graphs and bar graphs indicate mean with SEM error bars. Box plots contain 10^th^-90^th^ percentiles of data, with remaining data represented as individual points.

To examine neurological and behavioral phenotypes in DNMT3A^KO/+^ mice, we assessed basic measures of sensation and motor performance such as balance (ledge test, platform test), grip strength (inverted screen test), motor coordination (walking initiation, rotarod), and sensorimotor gating (pre-pulse inhibition). DNMT3A^KO/+^ mice were not significantly different in these assays (Figure S4A-G), indicating that heterozygous loss of DNMT3A does not grossly disrupt sensorimotor function. This allowed us to accurately assess more complex aspects of behavior and cognition.

We carried out a panel of assays with relevance to neuropathology observed in humans with DNMT3A mutations, including anxiety, autism, and intellectual disability. DNMT3A^KO/+^ mice displayed reduced exploratory behavior during open field testing, including reduced distance traveled and rearing (Figure 3C-D). DNMT3A^KO/+^ mice also displayed anxiety-like behaviors in this assay, as they spent less time in the center of the open field arena (Figure 3E). In tests of climbing behavior, DNMT3A^KO/+^ mice showed longer latency to climb to the bottom of a pole and to the top of mesh screens (Figure S4H-J), suggesting DNMT3A^KO/+^ mice display differences in volitional movement. To further assess anxiety-like behavior, we tested mice in the elevated plus maze and observed that DNMT3A^KO/+^ mice spent less time exploring the open arms of the maze with no change in percent entries made into all arms (Figure 3F, Figure S4K). Overall, these results demonstrate that the DNMT3A^KO/+^ mice display changes in exploratory behavior, suggesting altered emotionality and increased anxiety-like behaviors.

Analysis of common phenotypes examined in mouse models of autism (marble burying, three chamber social approach, ultrasonic vocalizations) revealed additional changes in behavior. We detected a significant reduction in marble burying activity for DNMT3A^KO/+^ mice, indicating alterations in repetitive digging behavior (Figure 3G). Testing of social interaction in the three chamber social approach test for adult mice (Yang et al., 2011) detected trends toward similar sociability and preference for social novelty in DNMT3A^KO/+^ compared to controls (Figure S4L). However, mutant mice showed reduced time investigating both mice and objects, as well as reduced overall activity (Figure 3H,I, Figure S4M). These results may further reflect the overall trend towards reduced exploration and anxiety-like phenotypes in these animals, instead of changes in sociability (Nygaard et al., 2019). Reduced maternal-isolation induced ultrasonic vocalizations (Barnes et al., 2017) were detected in DNMT3A^KO/+^ mice at postnatal day five, suggesting deficits in early pro-social behaviors or slight developmental delay in the normal acquisition of this behavior (Figure 3J). Together these results indicate alterations in behaviors commonly assessed in mouse models of autism (Chang et al., 2017; Simola and Granon, 2019; Takumi et al., 2019), with our findings suggesting a reduction in activity and exploration, as well as some changes in communication behaviors.

Intellectual disability is observed in patients with DNMT3A mutations, so we assessed learning and memory in the DNMT3A^KO/+^ mice using fear conditioning and Morris water maze tests. The mutant mice displayed largely similar recall performance to that of control mice in foot-shock induced fear conditioning, with similar levels of shock sensitivity (Figure 3K-M, Figure S4N). However, DNMT3A^KO/+^ mutants showed heightened freezing response during training, as well as contextual and auditory recall phases of conditioned fear testing (Figure 3K-M). Mutant mice also showed delayed extinction of freezing behavior in response to the auditory cue alone, which may indicate altered emotionality or cognition (see *methods* for discussion). Assessment of spatial and contextual memory by Morris water maze testing demonstrated that DNMT3A^KO/+^ mice were slower to learn to find a visible platform and did not learn the location of the hidden platform over time to the level of wild-type controls (Figure S4O-P). DNMT3A^KO/+^ mice also showed no differences in swimming speed compared to wild types (Figure S4Q-R). There were no significant effects on distance traveled in target zone or platform crossings in the probe trial, though DNMT3A^KO/+^ mice trended towards fewer platform crossings (Figure S4S,T). These findings suggest that DNMT3A^KO/+^ mutants do not show frank deficits in learning and memory but do display differences in task performance that further suggest altered emotionality or cognition in these mice. Our analyses demonstrate that heterozygous deletion of DNMT3A results in altered behavior in mice with relevance to anxiety and memory associated behaviors observed in patients with DNMT3A mutations. These data support a model in which DNMT3A haploinsufficiency can alter behavioral circuits to drive phenotypes in NDD.

### Global disruption of DNA methylation in the DNMT3A^KO/+^ brain

We next investigated the epigenomic defects that may underlie the altered behaviors observed in DNMT3A^KO/+^ mice. We first used sparse whole-genome bisulfite sequencing to efficiently survey effects on global DNA methylation levels for multiple brain regions and liver tissue isolated from wild type and DNMT3A^KO/+^ mice. This analysis detected limited reductions in genome-wide mCG levels in the DNMT3A^KO/+^ brain that were not apparent in the liver, a non-neural tissue (Figure 4A). In contrast, mCA levels were reduced by 30-50% across all brain regions examined in DNMT3A^KO/+^ mice (Figure 4B). DNA methylation across postnatal development in the cerebral cortex, the brain region with the highest levels of mCA at 8 weeks, suggests that deficits in mCA appear during initial accumulation of this methyl mark at 1-6 weeks (Figure 4B). Thus, global mCA levels in the brain appear to be highly sensitive to heterozygous DNMT3A disruption, while overall global mCG levels are largely maintained.

**Figure 4.**
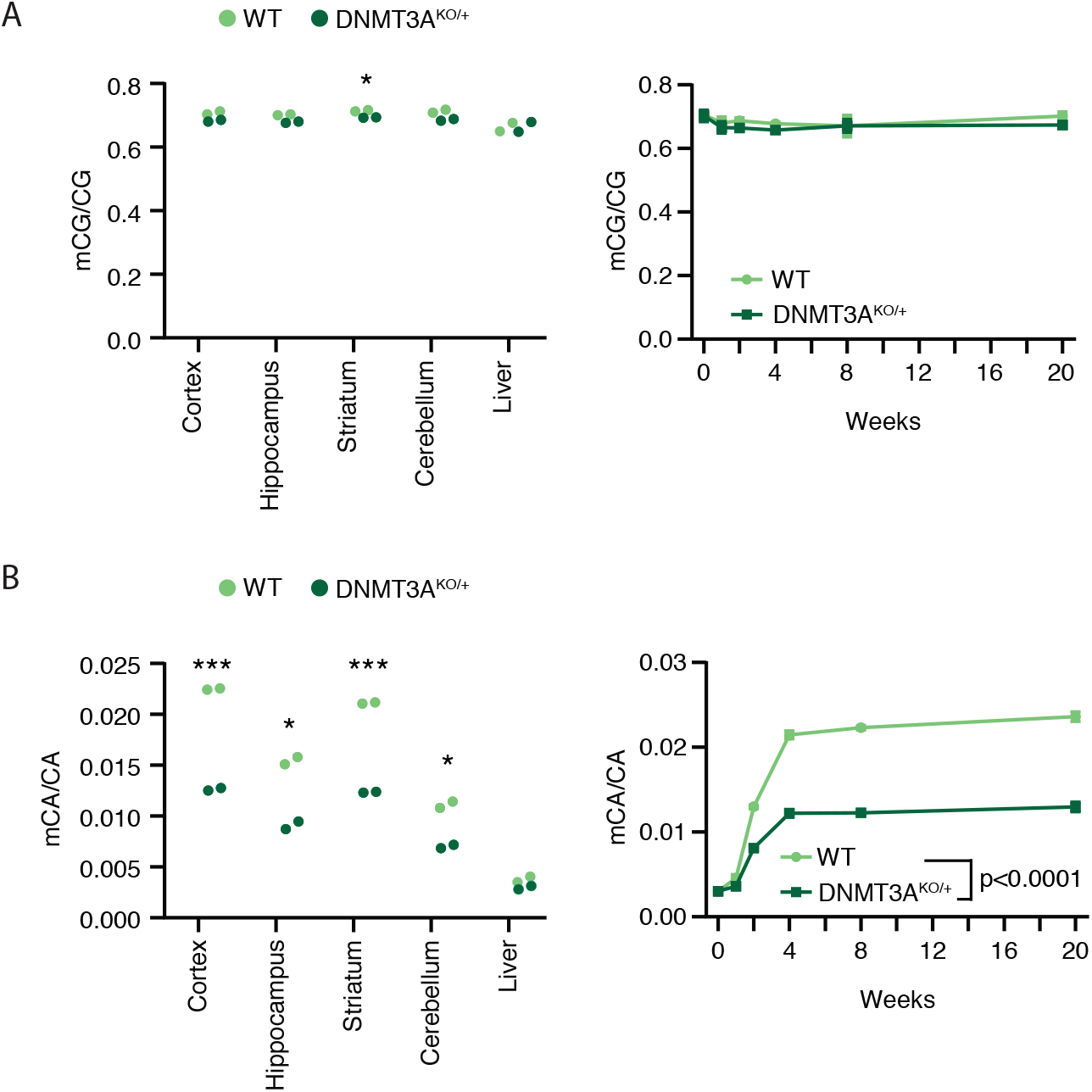
Global DNA methylation levels upon heterozygous loss of DNMT3A. (**A**) Global mCG levels in DNA isolated from tissues of 8-week old mice (left) (*, *P*<0.05; unpaired Student’s T-Test with Bonferroni correction), and developmental time course of global mCG (right), as measured by sparse whole genome bisulfite sequencing (WGBS). (**B**) Global mCA levels in DNA isolated from tissues of 8-week old mice (left) (***, *P*<0.001; *, *P*<0.05; unpaired Student’s T-Test with Bonferroni correction), and developmental time course of global mCA (right), as measured by sparse WGBS (*P*<0.0001 effect by genotype, F_(1,27)_=1024; *P*<0.0001 effect by age F_(5,27)_=884.6; n=3-4; two-way ANOVA). Line graphs indicate mean with SEM error bars.

DNA methylation at specific genomic elements, including promoters, enhancers, and gene bodies is thought to play an important role in regulating transcription. Alterations in methylation at these regions can impact gene expression to affect the development and function of the brain (Clemens et al., 2019; Nord and West, 2019; Stroud et al., 2017a). We therefore assessed changes in methylation at base-pair resolution by high-depth whole-genome bisulfite sequencing to identify potential changes in mCA and mCG at these important regulatory sites. For this analysis we focused on the cerebral cortex, as this region is enriched for mCA (Figure 4B) and is disrupted in ASD and MeCP2 disorders (Clemens et al., 2019; Satterstrom et al., 2019; Sceniak et al., 2016; de la Torre-Ubieta et al., 2016).

High-resolution analysis of mCG confirmed the subtle reduction in mCG across all classes of genomic elements (Figure 5A,D). We considered that CG dinucleotides in specific sites in the neuronal genome may be more sensitive to a partial reduction in DNMT3A activity. For example, in the hematopoietic system, heterozygous disruption of DNMT3A leads to reductions in DNA methylation in genomic regions that can be identified as sensitive to complete loss of DNMT3A (Cole et al., 2017). We therefore evaluated developmentally-regulated adult-specific CG-differentially methylated regions (CG-DMRs) previously identified in the cortex (Figure S5A) (Lister et al., 2013a). Because DNMT3A is the only *de novo* methyltransferase expressed in the postnatal brain, we hypothesized that adult-specific CG-DMRs might be sensitive to a reduction in enzyme activity. Indeed, we found that these sites build up mCG during postnatal development and do not become methylated in a brain-specific DNMT3A conditional knockout mouse (DNMT3A cKO) (Stroud et al., 2017a) (Figure S5B). Analysis of adult-specific CG-DMRs in the DNMT3A^KO/+^ model indicated that these sites are particularly sensitive to partial inactivation of DNMT3A compared to other regions genome wide (Figure 5A,C,D).

**Figure 5.**
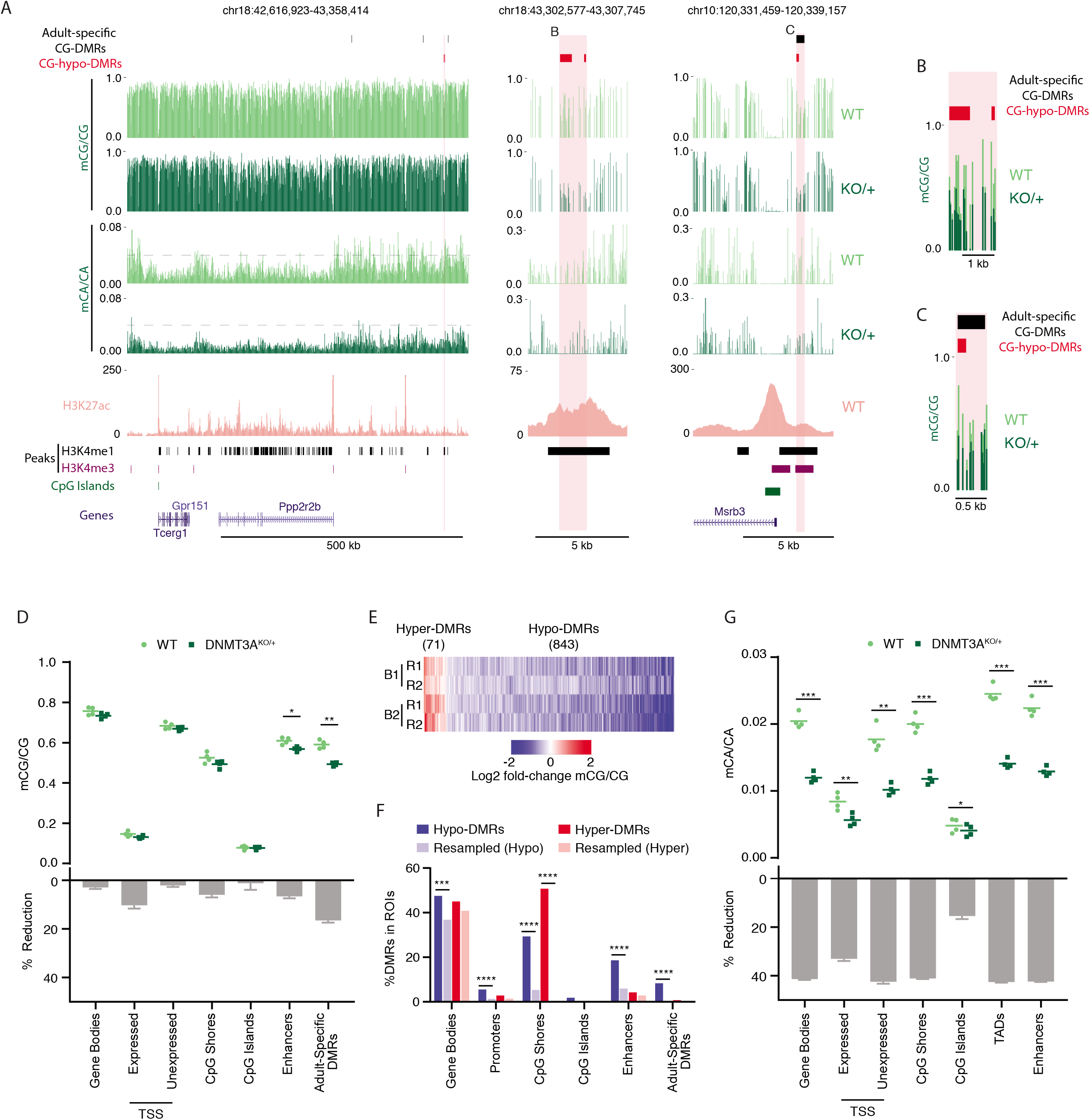
High-resolution analysis of DNA methylation changes in the DNMT3A^KO/+^ cerebral cortex. (**A**) Genome browser views of mCA and mCG in WT and DNMT3A^KO/+^ cerebral cortex as measured by high-depth WGBS. Broad view showing global reduction in mCA (left). Grey dashed line in mCA plots at 0.03 to facilitate visual comparison of global mCA levels between genotypes. Zoomed-in view of a DNMT3A^KO/+^ CG-hypo-DMR that overlaps an enhancer (center) and a DNMT3A^KO/+^ CG-hypo-DMR at a CpG-island shore that overlaps with an adult-specific DMR (right). WT H3K27ac ChIP-seq signal (Clemens et al., 2019), peaks of enhancer-associated H3K4me1 (Stamatoyannopoulos et al., 2012), peaks of promoter-associated H3K4me3 (Stamatoyannopoulos et al., 2012), CpG islands, and gene annotations (Haeussler et al., 2019) are shown below to illustrate overlap between DMRs and functional elements in the genome. (**B,C**) Overlay of mCG signal for DMR regions shown in **A**. (**D**) Mean mCG/CG level per replicate (top) and percent reduction, (DNMT3A^KO/+^-WT)/WT, (bottom) in WT and DNMT3A^KO/+^ cerebral cortex across indicated classes of genomic regions. Adult-specific DMRs were identified in the cortex (Lister et al., 2013a) (*, *P*<0.05; **, *P*<0.01; n=4 per genotype; paired Student’s T-Test with Bonferroni correction). (**E**) Heat map of CG DMRs called in the DNMT3A^KO/+^ cortex. Biological replicates (B1,B2) and technical replicates (R1,R2) are indicated. (**F**) Observed and expected (see *methods*) overlap between DNMT3A^KO/+^ cortex CG-DMRs and various genomic regions (****, *P*<0.0001; ***, *P*<0.001; Fisher’s Exact Test with Bonferroni correction). (**G**) Mean mCA/CA levels per replicate (top) and percent reduction, (DNMT3A^KO/+^-WT)/WT, (bottom) in WT and DNMT3A^KO/+^ cortex across indicated classes of genomic regions (*, *P*<0.05; **, *P*<0.01; ***, *P*<0.001; n=4; paired Student’s T-Test with Bonferroni correction). Box plots indicate median and quartiles.

To further search for local sites of altered mCG in the DNMT3A^KO/+^, we performed *de novo* calling of mCG differentially methylated regions using the BSmooth algorithm (Hansen et al., 2012). We identified 843 hypo- and 71 hyper-CG-DMRs across the genome that met high stringency filters for size and reproducibility (Figure 5A-C,E, Figure S5C, see *methods*). These hypo-DMRs significantly overlap with the previously identified adult-specific CG-DMRs (Lister et al., 2013a) (Figure 5F), further supporting the idea that DNMT3A is haploinsufficient for postnatal mCG deposition at these sites. Examination of the genomic distribution of all DNMT3A^KO/+^ CG-DMRs revealed significant overlap of hypo-DMRs with putative enhancer regions, gene bodies, and promoters (Figure 5A-C,F). DMRs were also highly enriched for overlap with CpG island shores, regions disrupted in studies of DNMT3A mutation outside of the nervous system (Cole et al., 2017; Spencer et al., 2017) (Figure 5F). Because a substantial percentage of mCG in neurons can occur in an oxidized, hydroxymethyl form (hmCG), we further performed oxidative bisulfite sequencing analysis of DNA from the cortex. This analysis revealed no clear evidence of differential effects on the oxidized or unoxidized forms of mCG across the genome in the DNMT3A^KO/+^ (Figure S5D). Together these findings indicate that a small subset of mCG sites are particularly sensitive to heterozygous loss of DNMT3A. The localization of these CG-DMRs to regulatory elements suggests that these methylation changes could impact gene expression.

We next examined the profile of mCA at higher resolution, assessing genomic elements of different scales that have relevance to gene regulation. In contrast to the limited mCG changes in the DNMT3A^KO/+^, analysis of mCA levels detected consistent 30-50% reductions at nearly all genomic regions examined (Figure 5G). This was true of gene bodies, promoters, and CpG island shores. CpG island sites, which show very low mCA levels in wild-type cortex, displayed less reduction of mCA, possibly due to floor effects in bisulfite-sequencing (see *methods)*. Comparison of mCA changes in each class of genomic elements as a function of wild-type mCA levels suggested that consistent reductions occurred across the genome independent of the normal levels of mCA (Figure S5E). This suggests that changes in mCA levels in the DNMT3A^KO/+^ do not preferentially impact specific classes of genomic elements or become substantially more severe in some regions based on the level of mCA that normally is deposited.

Recent analysis has demonstrated that topologically-associating domains (TADs) of chromatin folding are regions of organization for mCA that can impact gene regulation (Clemens et al., 2019). The “set-point” level of mCA within TADs is associated with the level of mCA at enhancers within TADs and high-mCA enhancers found in high-mCA TADs are particularly robust targets of repression by MeCP2 (Clemens et al., 2019). We therefore specifically assessed mCA levels at TADs and enhancers genome-wide. This analysis detected reductions in TAD mCA levels that were similar to global reductions in mCA at other genomic elements (Figure 5G). Enhancers also showed this pervasive depletion of mCA (Figure 5G). Like other genomic elements, these effects were consistent for TADs and enhancers with differing wild-type levels of mCA (Figure S5E). Thus, widespread loss of mCA for TADs and enhancer elements occurs in DNMT3A^KO/+^ mice and has the potential to impact epigenetic control of regulatory elements by MeCP2.

### Enhancer dysregulation results from methylation deficits in DNMT3A^KO/+^ mice

We next examined how disruption of DNA methylation can affect epigenetic regulation in DNMT3A^KO/+^ neurons to alter gene expression and disrupt nervous system function. Recent analysis indicates that mCA serves as a binding site for MeCP2 to mediate neuron-specific gene regulation, in part by controlling the activity of distal regulatory enhancer elements (Clemens et al., 2019). Loss of MeCP2 in mice leads to genome-wide upregulation of the activating mark Histone H3 lysine 27 acetylation (H3K27ac) at enhancers that contain high levels of mCA and mCG sites, while overexpression of MeCP2 leads to reciprocal downregulation of highly methylated sites. Alterations in enhancer activity in MeCP2 mutants are linked to dysregulation of genes that can then drive nervous system dysfunction. These findings suggest that reduced CA methylation in the DNMT3A^KO/+^ would remove binding sites for MeCP2 within enhancers. This mCA reduction could then result in dysregulation of enhancer activity that partially phenocopies the effects we have observed in MeCP2 mutant mice.

To investigate this possibility directly, we quantified the change in mCA binding sites in the DNMT3A^KO/+^ for enhancers significantly repressed by MeCP2 (Clemens et al., 2019). These enhancers contain a large number of mCA sites due to high mCA/CA levels and an enrichment of CA dinucleotides within these sequences. As a result, we found that the global 30-50% reduction of mCA in the DNMT3A^KO/+^ leads to a larger loss in the total number of mCA sites at MeCP2-repressed enhancers than at other enhancers genome-wide (Figure 6A,B). Thus MeCP2-repressed enhancers are particularly susceptible to mCA binding site loss from heterozygous mutation of DNMT3A.

**Figure 6.**
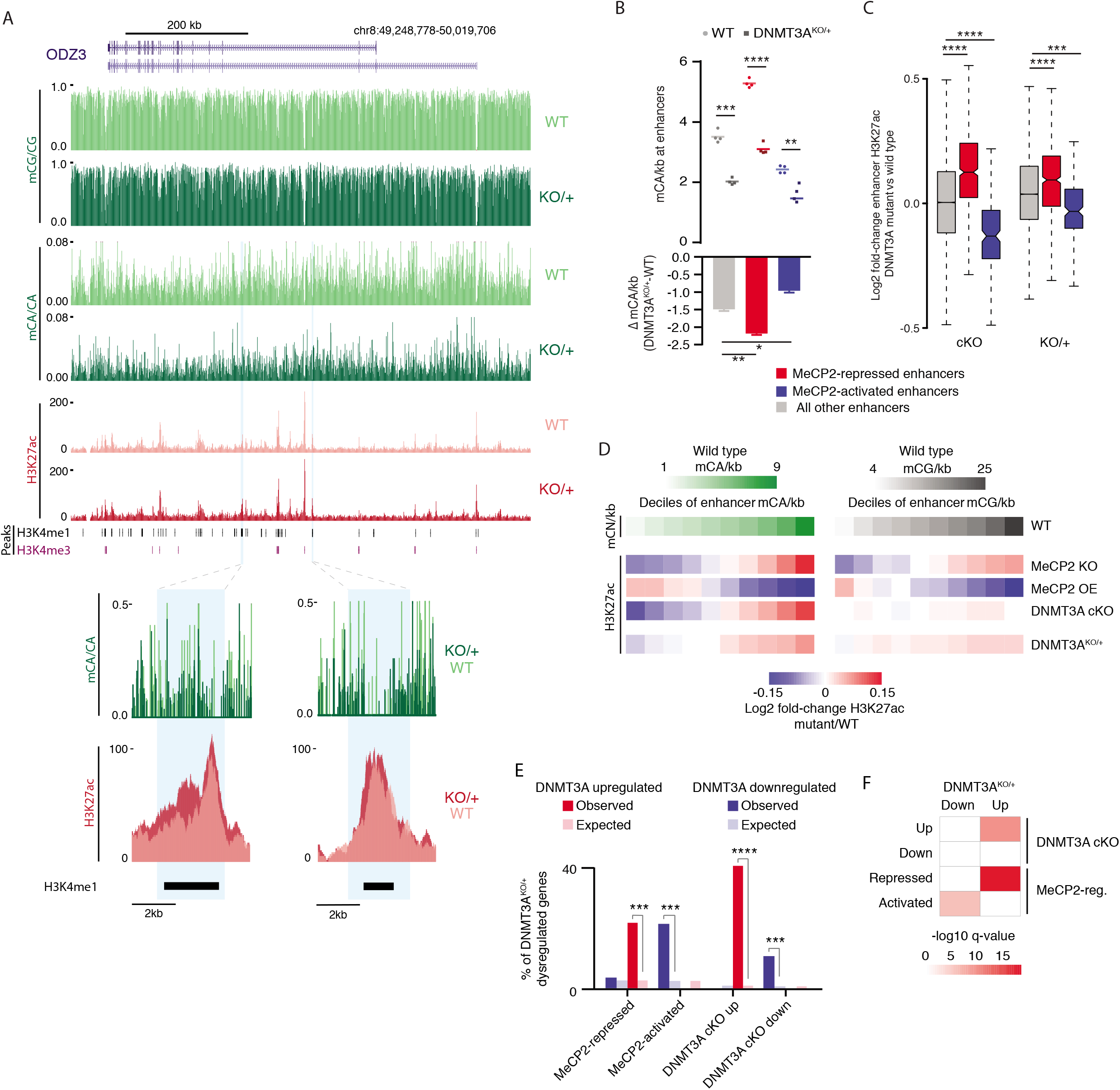
DNMT3A^KO/+^ enhancer dysregulation and transcriptomic pathology overlaps with MeCP2 mutants. (**A**) Genome browser view of DNA methylation and H3K27ac ChIP-seq data from WT and DNMT3A^KO/+^ cerebral cortex (top). Overlaid H3K27ac signal and mCA/CA levels at enhancer regions highlighted in blue that were identified as dysregulated enhancers upon disruption of mCA or MeCP2 (Clemens et al., 2019) (bottom). (**B**) Mean mCA sites/kb in WT and DNMT3A^KO/+^ cortex (top) and number of mCA sites/kb lost in the DNMT3A^KO/+^ cortex (bottom) for enhancers significantly dysregulated in MeCP2 mutants (*, *P*<0.05; **, *P*<0.01; ***, *P*<0.001; ****, *P*<0.0001; n=4; paired Student’s T-Test with Bonferroni correction). (**C**) Boxplot of fold-change in H3K27ac signal in DNMT3A cKO and the DNMT3A^KO/+^ cortex for enhancers defined as significantly dysregulated in MeCP2 mutants. (***, *P*<10^-8^; **** *P*<10^-12^; n=5 biological replicates of DNMT3A^KO/+^ and WT; Wilcoxon test) (**D**) Heatmap of changes in H3K27ac signal indicated mutants across deciles of enhancers sorted by wild-type mCA or mCG sites (*P*<2.2e-16, DNMT3A^KO/+^ mCA/kb; *P*<2.2e-16, DNMT3A^KO/+^ mCG/kb; Spearman Rho correlation). (**E**) Observed versus expected overlap of significantly dysregulated genes (padj. < 0.1) in the DNMT3A^KO/+^ and genes dysregulated in DNMT3A cKO or MeCP2 mutant mice (***, *P*<1e-5; ****, *P*<1e-10; hypergeometric test). (**F**) Significance of gene set expression changes in the indicated direction in the DNMT3A^KO/+^ cortex for GAGE analysis of gene sets identified as dysregulated in DNMT3A cKO or MeCP2 mutant mice (Clemens et al., 2019). Note: legend is shared in **B** and **C.** Box plots indicate median and quartiles.

To determine if the reduction of mCA sites at MeCP2-repressed enhancers affects their activity, we assessed changes in enhancer activation level by H3K27ac ChIP-seq analysis of the DNMT3A^KO/+^ and wild-type cerebral cortex. This analysis revealed significant changes in acetylation at MeCP2-repressed enhancers (Figure 6A,C). Consistent with these effects arising from 30-50% loss of the mCA that normally builds up post-mitotically at enhancers, we detect changes that are concordant with, but smaller than, those caused by complete loss of postmitotic mCA in the DNMT3A cKO (Clemens et al., 2019) (Figure 6C).

Although significantly dysregulated enhancers can be detected in MeCP2 mutants, broad sub-significance-threshold effects also occur genome-wide upon MeCP2 mutation, with nearly all enhancers across the genome undergoing dysregulation that is proportional to the number of mC binding sites at these regions (Figure 6D) (Clemens et al., 2019). Analysis of H3K27ac changes at enhancers based on the normal density of mCA sites in these sequences genome-wide revealed broad mCA-associated derepression of enhancers in DNMT3A^KO/+^ cortex that is similar to, but smaller in magnitude than, the effects observed in DNMT3A cKO and MeCP2 knockout mice (MeCP2 KO). These effects are also reciprocal to effects observed in MeCP2 overexpression mice (MeCP2 OE). Consistent with the limited disruption of mCG genome wide in the DNMT3A^KO/+^ mice, there was more limited association between changes in enhancer activity and the level of mCG at these sequences. This contrasts with MeCP2 mutants in which loss of protein binding at both mCG and mCA sites leads to enhancer dysregulation that is associated with both mCA and mCG (Clemens et al., 2019) (Figure 6D). Together, these findings demonstrate that loss of half of the normal mCA sites in the DNMT3A^KO/+^ cortex results in enhancer dysregulation that overlaps with MeCP2 mutant mice, uncovering a role for shared neuronal chromatin pathology between DNMT3A and MeCP2 disorders.

### Overlapping transcriptional pathology between DNMT3A^KO/+^, MeCP2 disorders, and ASD

The epigenetic alterations we observe in DNMT3A^KO/+^ cerebral cortex can have direct consequences on gene expression to drive neurological dysfunction in mice. Furthermore, the overlapping effects on enhancers that we observe between DNMT3A^KO/+^ and MeCP2 mutant mice suggests that there may be shared transcriptional pathology occurring upon loss of mCA in DNMT3A disorders and through disruption of MeCP2 in Rett syndrome and MeCP2-duplication syndrome. We therefore assessed changes in gene expression in DNMT3A^KO/+^ mice, interrogating the extent to which these effects overlap with those observed in MeCP2 mutants and upon complete disruption of mCA in the DNMT3A cKO. RNA-seq of DNMT3A^KO/+^ cerebral cortex identified subtle changes in gene expression that are consistent in magnitude with effects observed in other heterozygous NDD models (Fazel Darbandi et al., 2018; Gompers et al., 2017; Katayama et al., 2016) (Figure S6A). Gene set enrichment analysis on differential expression data revealed dysregulation of Gene Ontology terms relating to neuronal development and function (Figure S6B), suggesting that alterations in gene expression resulting from DNMT3A heterozygous disruption could drive the behavioral alterations that we observed.

While a limited gene set is detected as significantly dysregulated in the DNMT3A^KO/+^, we considered if genome-wide alterations in enhancer activity could lead to wide-spread, subtle dysregulation of gene expression that is below the threshold of detection for individual genes. In this way, the transcriptional pathology in the DNMT3A^KO/+^ brain could overlap with the subthreshold genome-wide effects observed upon loss of neuronal mCA (DNMT3A cKO) and in models of Rett syndrome (MeCP2 KO) and ASD (MeCP2 OE) (Clemens et al., 2019; Gabel et al., 2015). Indeed, the significantly dysregulated genes in the DNMT3A^KO/+^ overlapped extensively with genes identified as significantly dysregulated in DNMT3A cKO and MeCP2 mutant mice (Clemens et al., 2019), supporting the notion of shared gene expression effects between these mouse models (Figure 6E). To more comprehensively assess the degree to which transcriptome-wide changes in the DNMT3A^KO/+^ phenocopy these MeCP2 mutant and DNMT3A cKO models, we performed Generally Applicable Gene-set Enrichment (GAGE) analysis (Luo et al., 2009) of all genes detected as dysregulated in these mutant models. This revealed highly significant, concordant changes in gene expression in the DNMT3A^KO/+^ for dysregulated gene sets detected upon loss of mCA in the DNMT3A cKO and in MeCP2 mutant models (Figure 6F).

Having detected overlap in transcriptomic pathology between models of DNMT3A and MeCP2 disorders, we sought to explore if shared gene expression signatures in the DNMT3A^KO/+^ mice extend to models of disorders that do not have as clear mechanistic links to DNMT3A disorders. We therefore tested if DNMT3A^KO/+^ mice show significant alterations in gene sets identified as dysregulated in other mouse models of NDD and human gene sets implicated as altered in the autistic brain. GAGE analysis across multiple datasets detected highly significant dysregulation of gene sets identified in CHD8 and PTEN mouse models of overgrowth and ASD (Gompers et al., 2017; Katayama et al., 2016; Tilot et al., 2016) as well as the SetD5 model of NDD (Sessa et al., 2019) (Figure 7A). These findings support a role for overlapping gene dysregulation underlying common symptomology found in affected individuals carrying mutations in distinct genes.

**Figure 7.**
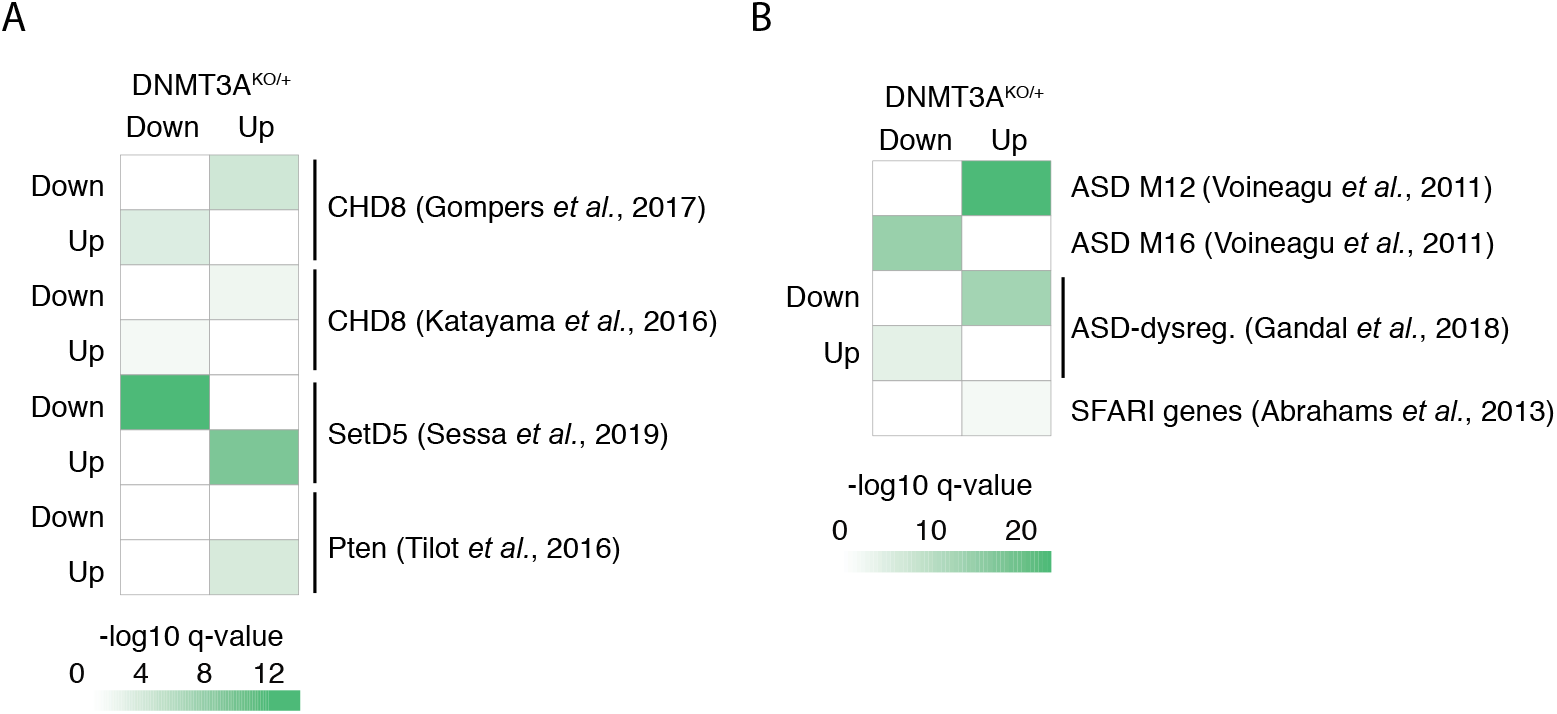
Gene dysregulation in the DNMT3A^KO/+^ overlaps with other ASD/NDD disorders. (**A**) GAGE analysis of expression changes in DNMT3A^KO/+^ for dysregulated gene sets identified in studies of NDD mouse models (Gompers et al., 2017; Katayama et al., 2016; Sessa et al., 2019; Tilot et al., 2016) (n=7 biological replicates of DNMT3A^KO/+^ and WT). (**B**) GAGE analysis of expression changes in DNMT3A^KO/+^ for gene sets identified in studies of human ASD. Gene sets: ASD module 12 (synaptic) and 16 (immune) identified in weighted-gene coexpression analysis of human ASD brain (Voineagu et al., 2011), and ASD-dysregulated genes previously identified (Abrahams et al., 2013; Gandal et al., 2018).

Analysis of human gene sets detected as dysregulated in ASD postmortem brains (Gandal et al., 2018; Voineagu et al., 2011) showed significant changes in the DNMT3A^KO/+^ cortex (Figure 7B). This analysis also indicated upregulation of candidate genes linked to ASD from human genetics studies (Abrahams et al., 2013; Banerjee-Basu and Packer, 2010) (Figure 7B). In addition, analysis of co-expression modules of human brain development (Parikshak et al., 2013) showed overlap with several neurodevelopmental modules including those that increase during early cortical development and are enriched for ASD risk genes (M13, M16, and M17) (Figure S6C). Modules involved in regulation of nucleic acids and gene regulation that are expressed early in development and decrease over time are also increased upon heterozygous loss of DNMT3A (M2 and M3) (Figure S6C). These results indicate that important sets of genes with opposing developmental trajectories and function are altered upon loss of DNMT3A regulation. Notably, control resampling analysis indicated that significant dysregulation of these mouse and human gene sets was not driven by enriched expression of these genes in the cortex (Figure S6D). Together these findings suggest that the DNMT3A^KO/+^ mouse shares overlapping transcriptional pathology with gene expression changes underlying ASD.

## Discussion

Our functional analysis of NDD-associated DNMT3A mutations together with our *in vivo* studies provide an initial working model of molecular etiology in DNMT3A disorders. Diverse *de novo* missense mutations that arise in affected individuals disrupt enzyme function by disabling the capacity of the enzyme to localize to chromatin in the nucleus, altering the ADD-regulatory domain, or disrupting the activity of the methyltransferase domain. Loss-of-function effects resulting from these missense mutations, as well as early truncations or gene deletions, lead to insufficient DNMT3A activity. This causes deficits in deposition of mCG at specific sites during development and a massive deficit in postnatal mCA accumulation throughout the brain. These changes in DNA methylation lead to alterations in epigenomic regulation, including subtle but wide-spread disruption of mCA-MeCP2-mediated enhancer regulation in adult neurons, resulting in gene expression changes that can drive deficits in nervous system function.

Our studies of DNMT3A mutations not only provide insight into the molecular etiology of DNMT3A disorders, but also serve as a model for understanding the functional effects of diverse *de novo* mutations underlying neurodevelopmental disorders. Exome sequencing studies have identified a large and growing list of mutations in genes encoding epigenetic regulators in individuals with NDD. Many of these are missense mutations and occur as heterozygous disruptions (McRae et al., 2017; Satterstrom et al., 2019), leaving it unclear if simple loss-of-function effects are sufficient to drive pathology through haploinsufficiency, or if more complex effects play a role when individual amino acids are altered. In addition, while identification of multiple mutations in a gene can implicate disruption of the gene as causative for NDD, it remains possible that a subset of the mutations identified in affected individuals, particularly missense mutations, are not in fact deleterious or causative. Functional testing of these variants is therefore necessary to determine if they may underlie disease. Here, our analysis of DNMT3A mutations in multiple functional assays has uncovered diverse mechanisms by which the protein can be disrupted while pointing to a shared loss of function in the deposition of neuronal DNA methylation. Notably, it is only by assessing multiple aspects of protein function (i.e. expression, localization, activity, and cellular mCA levels) that we can detect deficits for each mutation tested. For example, mutation of the ADD domain disrupts deposition of mCA, possibly due to loss of regulation that can only be assessed in the endogenous chromatin context. Together our findings establish the deleterious effects of diverse DNMT3A mutations and underscore the importance of multidimensional analysis of *de novo* mutations to fully assess their potential role in NDD.

Our *in vivo* analyses show that heterozygous deletion of DNMT3A mirrors multiple key features of DNMT3A disorders, including tall stature (increased long bone length), increased body weight, and behavioral alterations. Detection of robust anxiety-like phenotypes in multiple assays, deficits in pro-social communication, and alterations in repetitive behaviors align with observed human phenotypes. In contrast, lack of strong deficits in learning and memory assays in our mouse model may indicate that some regions and systems in humans are more susceptible to DNMT3A disruption than in mice. However, we do detect alterations in behavior in these assays (Figure 3K-M, Figure S4O-T) and the lack of strong deficits may also reflect insensitivity of the methods used to measure specific aspects of disrupted cognition. In all, our *in vivo* analysis indicates that heterozygous deletion of DNMT3A results in effects which can guide future studies of molecular, cellular, and organismal dysfunction caused by mutation of DNMT3A.

We employed the DNMT3A^KO/+^ mouse experimental system to assess how heterozygous DNMT3A disruption impacts epigenetic regulation in the brain. Our analysis of DNA methylation in tissues from DNMT3A^KO/+^ mice detected very subtle changes in genome-wide mCG levels across brain regions, with no global mCG effects in non-neural tissue (Figure 4A). Analysis of local changes in mCG in the brain detected evidence of disrupted CG methylation at sites methylated during postnatal development (i.e. adult hyper CG-DMRs). In addition, multiple hypo-CG-DMRs can be detected at regulatory elements including enhancers. While limited, these effects have the potential to alter gene expression and contribute to neurological alterations in these mice. The limited nature of mCG effects is likely due to the redundant function of the other DNA methyltransferases. The maintenance methyltransferase DNMT1 has the capacity to preserve existing mCG patterns during cell divisions (Jeltsch et al., 2018). In addition, the *de novo* methyltransferase DNMT3B is expressed with DNMT3A in many tissues during early development and could provide critical redundancy for mCG patterning (Okano et al., 1999). Nonetheless, the site-specific changes in mCG are also likely to occur in early development and in non-neural tissues. For example, constitutive heterozygous deletion of DNMT3A has been shown to disrupt mCG patterns in the blood and alter hematopoietic lineages (Cole et al., 2017). These changes in mCG may contribute to changes in growth and other phenotypes observed in mice and humans.

In contrast to mCG, we detect a global reduction in mCA to approximately 30-50% of wildtype levels in DNMT3A^KO/+^ cortex, striatum, cerebellum, and hippocampus (Figure 4B). These results generalize and extend findings in the hypothalamus (Sendžikaitė et al., 2019), demonstrating the susceptibility of broad neuronal types and circuits to heterozygous loss of DNMT3A. The susceptibility of mCA to heterozygous loss of DNMT3A is likely due to several related factors. For example, DNMT3B is not expressed in postnatal neurons (Lister et al., 2013a), and DNMT1 is not capable of depositing mCA (Jeltsch et al., 2018), making all mCA build-up in neurons dependent on DNMT3A. In addition, the enzyme has slow kinetics for activity on CA sites (Zhang et al., 2018) and deposition of mCA genome-wide by DNMT3A must take place in a restricted time window (1-6 weeks) when the enzyme is highly expressed and active in neurons (Clemens et al., 2019; Lister et al., 2013a; Stroud et al., 2017a). These constraints may make enzyme levels limiting for mCA accumulation in neurons, providing an explanation for why global mCA in the brain is sensitive to DNMT3A gene dosage. Notably, our findings suggest that manipulations that activate the remaining DNMT3A, or prolong its high early postnatal expression, might rescue deficits in mCA deposition. Conversely, duplication of the DNMT3A gene could result in too much deposition of mCA and possibly cause significant neural dysfunction akin to those effects seen in MeCP2 duplication disorder. Future studies can assess the feasibility of rescue approaches and explore if DNMT3A duplication alters brain function.

Our analysis of chromatin changes downstream of altered DNA methylation has uncovered a striking point of shared molecular disruption across models of DNMT3A disorders, Rett syndrome, and MeCP2 duplication syndrome. While the clinical profile and pathophysiology of DNMT3A disorders is clearly distinct from MeCP2 disorders, we have shown here that loss of approximately a quarter of MeCP2 binding sites across the neuronal genome in the DNMT3A^KO/+^ cortex results in subtle but wide-spread disruption of mCA-associated enhancer regulation that partially phenocopies loss of MeCP2. This enhancer dysregulation can be linked to shared alterations in gene expression across these models (Clemens et al., 2019) (Figure 6). Given the critical roles of MeCP2-regulated genes for nervous system function (Gabel et al., 2015; Lagger et al., 2017; Lyst and Bird, 2015), these epigenomic and transcriptomic effects likely contribute to aspects of neurologic dysfunction observed in DNMT3A disorders. The persistence of many mCA and mCG binding sites for MeCP2 in the DNMT3A^KO/+^ may partially explain how DNMT3A mutations manifest with less severe symptomology than in Rett Syndrome. In addition, absence of DNMT3A early in prenatal development can contribute to overgrowth and other non-overlapping aspects of DNMT3A and MeCP2 disorders. Together, our findings show that disruption of mCA-MeCP2 mediated enhancer regulation likely contributes to three disorders with distinct symptomology, defining a site of convergent molecular etiology underlying heterogeneous clinical syndromes.

Our transcriptomic analysis of changes of ASD/NDD gene sets in DNMT3A mice has further detected overlap with NDD beyond MeCP2 disorders, including both mouse models of NDD/ASD (CHD8) and gene sets identified in human idiopathic ASD. As additional transcriptomic studies of mouse models and human NDD brain emerge, systematic analyses of gene expression effects can identify shared aspects of transcriptional pathology that can contribute to cognitive and social deficits across diverse causes of NDD. Notably, the large number of chromatin modifying enzymes mutated in these disorders raises the possibility that shared transcriptomic effects emerge from common chromatin pathology. Our study has identified alterations in mCA and enhancer regulation as a potential site of convergent dysfunction in MeCP2 and DNMT3A disorders. Future studies may identify additional gene disruptions in which alterations in mCA and enhancer dysregulation contribute to molecular pathology, expanding the role of “methylopathies” in neurodevelopmental disease.

## Acknowledgements

We thank the Division of Comparative Medicine at Washington University in Saint Louis for their assistance with mouse husbandry and veterinary support. We are grateful for sequencing support from the Genome Technology Access Center and the Center for Genome Sciences and Systems Biology Spike in Cooperative at Washington University in Saint Louis. We thank members of the Dougherty lab including S. Maloney, K. McCullough, and M. Rieger for assistance in USV experiments. We thank J. Edwards, J. Goodman, and J. Yi for critical feedback on the experimental design and manuscript. We thank A. Smith and T. Ley for helpful discussions. This work was supported by NIH 5T32GM00815133 to D.L.C, by NIH 5T32GM007067 and F31NS108574 to A.W.C., and by the Klingenstein-Simons Fellowship Fund, the G. Harold and Leila Y. Mathers Foundation, the Brain and Behavior Research Foundation, the Simons Foundation for Autism Research Initiative, and NIMH R01MH117405 to H.W.G.

## Author Contributions

D.L.C and D.Y.W. are joint first authors, as each led critical components of the project and analysis. D.L.C., J.R.Ma., and Y.R.L. generated and analyzed *in vitro* biochemical data. Y.R.L. and S.A.N. generated primary neuronal culture samples. D.L.C. and J.R.Ma. generated skeletal samples. N.M.K. and C.A.H. carried out craniofacial analysis and D.L.C. carried out long bone analysis. D.L.C., J.R.Ma., D.F.W., and J.D.D. carried out behavioral tests and analysis. D.L.C., J.R.Ma., J.R.Mo., Y.R.L., and A.W.C. generated genomic data. D.Y.W. developed analysis algorithms and pipelines. D.L.C., D.Y.W., J.R.Mo, and A.W.C. completing genomic analyses. H.W.G. conceived the project and H.W.G., D.L.C., and D.Y.W. designed the experiments. H.W.G., D.L.C., and D.Y.W. wrote the manuscript and all authors contributed to manuscript editing and revisions.

---

**Figure S1.**
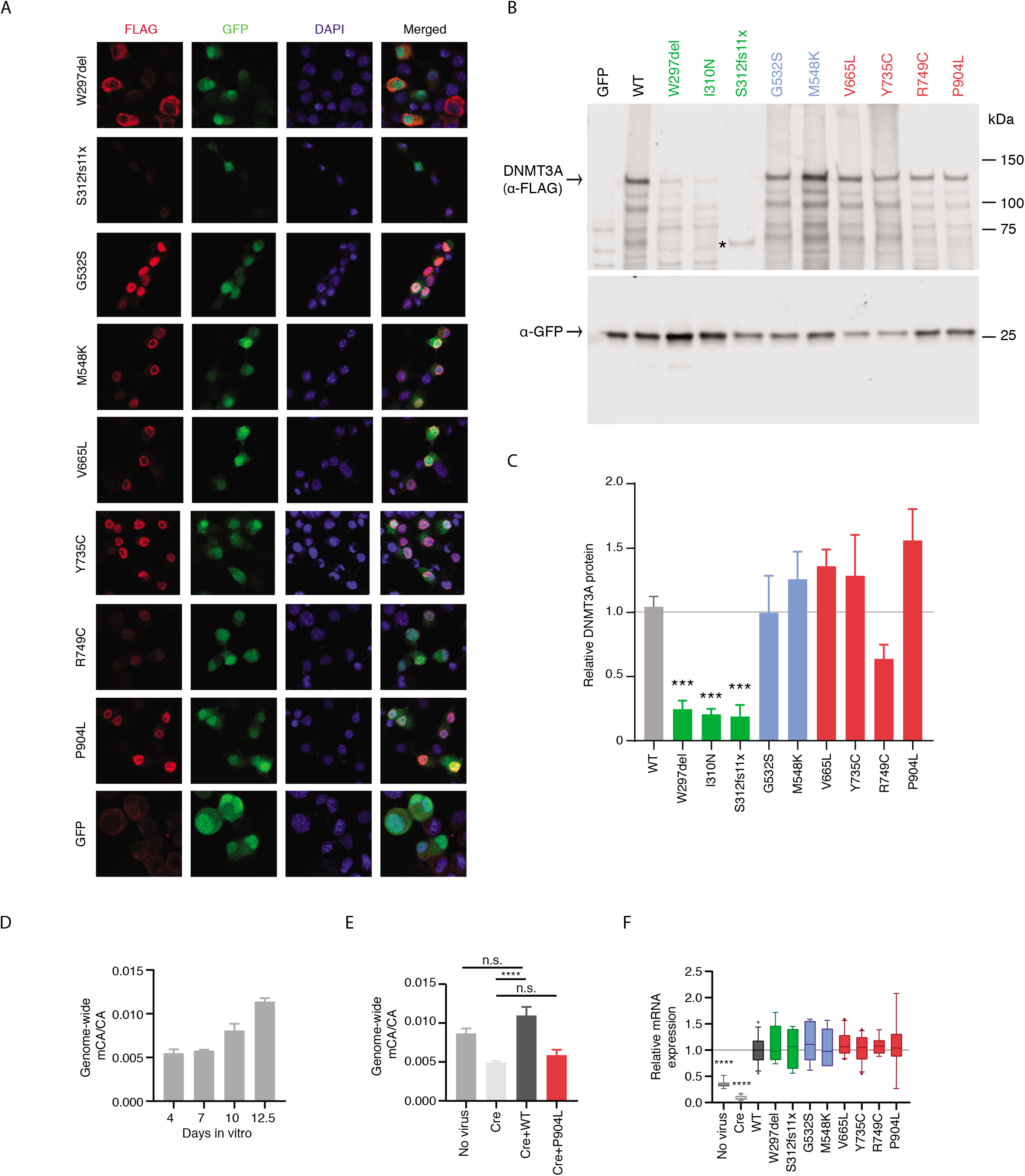
Related to Figures 1 and 2. (**A**) Example images of DNMT3A mutant protein immunocytochemistry. Scale bar = 20μm. (**B**) Full example immunoblot from Figure 1b with the truncated S312fs11x DNMT3A mutant protein indicated with an asterisk. (**C**) Quantification of immunoblot signal of DNMT3A (α-FLAG) from mutant proteins (***, *P*<0.001; n=6-30; unpaired Student’s T-Test with Bonferroni correction). (**D**) Genome wide mCA levels over time in neuronal cortical cultures, as measured by sparse WGBS (*P*=0.0035 effect by time, F_(3,4)_=29.45, n=2; one-way ANOVA). (**E**) Genome-wide mCA levels in DNMT3A mutant add-back cortical cultures (****, *P*<0.0001; *, *P*<0.05; n=7-11; planned unpaired Student’s T-Tests with Bonferroni correction). (**F**) qRT-PCR of *Dnmt3a* normalized to *Actb* for samples chosen for WGBS. (****, *P*<0.0001; n=4-11; one-sample Student’s T-Test with Bonferroni correction). Bar graphs indicate mean with SEM error bars.

**Figure S2.**
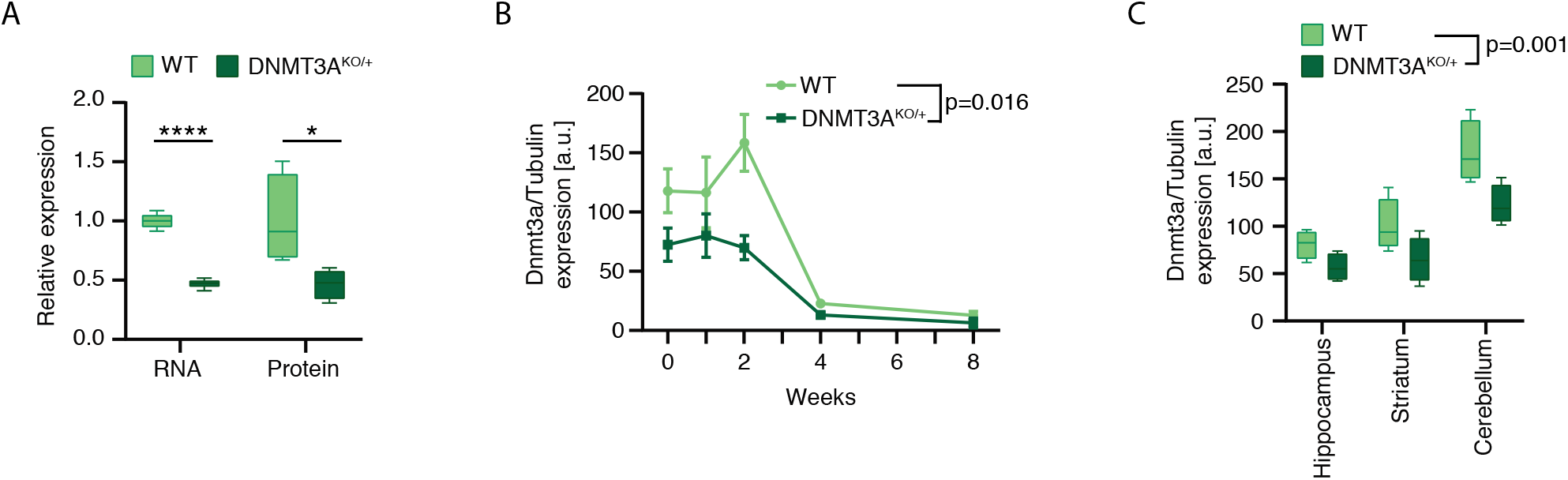
Related to Figures 3, 4, 5, 6 and 7. (**A**) Normalized *Dnmt3a* mRNA and protein expression from 2-week cortices of DNMT3A^KO/+^ and wild-type littermates (****, *P*<0.0001; *, *P*<0.05; mRNA n=5, protein n=4, unpaired Student’s T-Test). (**B**) Protein expression of DNMT3A normalized to α-Tubulin measured by western blotting for cerebral cortex of DNMT3A^KO/+^ and wild-type littermates over developmental time (*P*=0.0157 effect by genotype, F_(1,16)_=7.303, n=2-5; two-way ANOVA). (**C**) Protein expression of DNMT3A normalized to α-Tubulin measured by western blotting for hippocampus, striatum, and cerebellum (*P*=0.0010 effect by genotype, F_(1,18)_=15.48, n=4; two-way ANOVA). Line plots indicate mean with SEM error bars.

**Figure S3.**
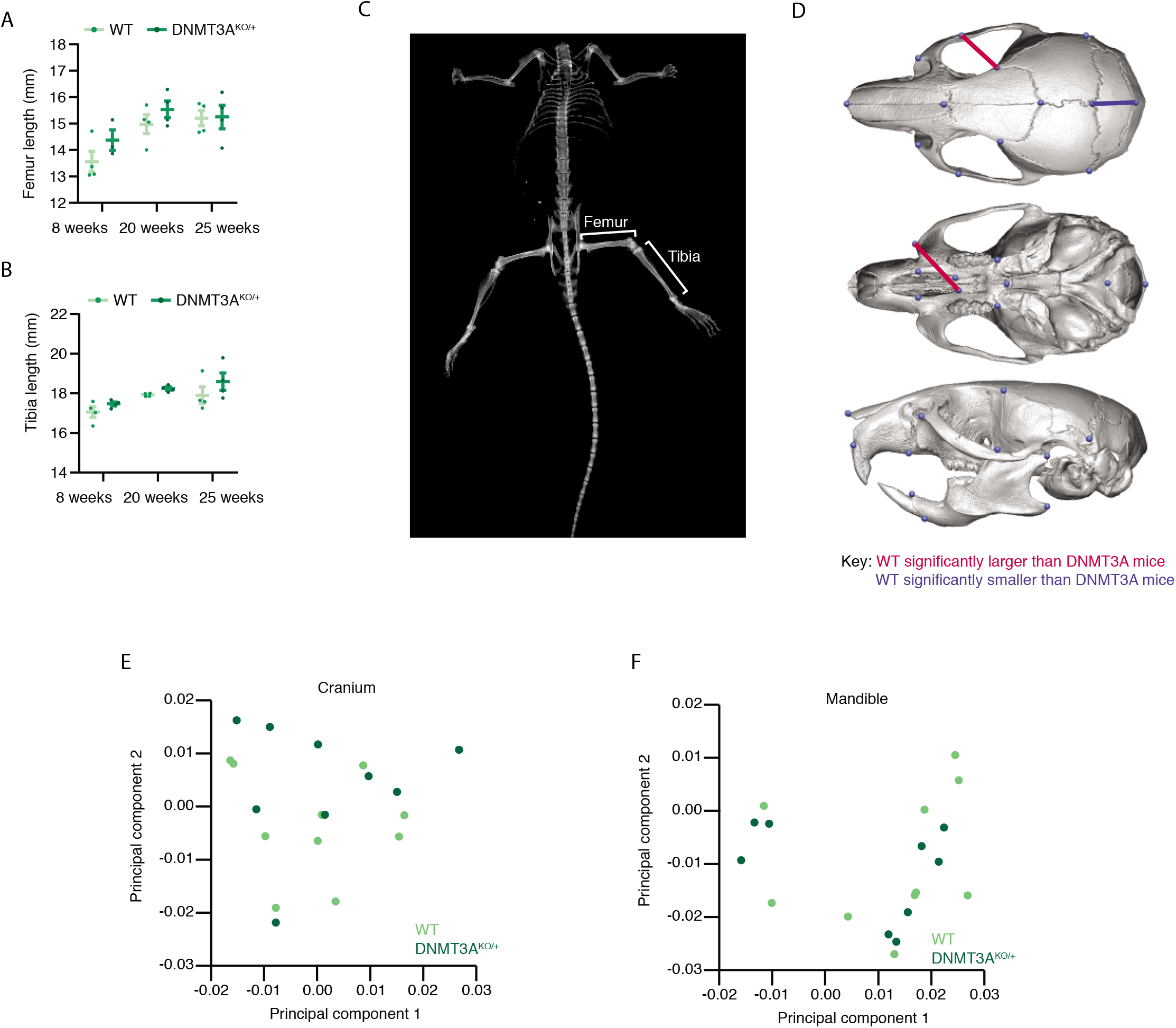
Related to Figure 3. (**A,B**) Measurements of (**A**) femur and (**B**) tibia by dual x-ray imaging in WT and DNMT3A^KO/+^ mice at 8, 20, and 25 weeks of age. Lines indicate mean with SEM error bars. (**C**) Example dual x-ray image of mouse body with femur and tibia indicated. (**D**) Example of reconstructed skull from μCT imaging with landmarks used for craniofacial analysis shown. Red line indicates distance that is significantly larger in the WT compared to the DNMT3A^KO/+^, while blue line indicates distance that is significantly smaller in the WT compared to the DNMT3A^KO/+^ (*P*<0.05). (**E,F**) Principal component analysis of (**E**) cranial and (**F**) mandibular shape shows no clear separation between groups along PC1 or PC2.

**Figure S4.**
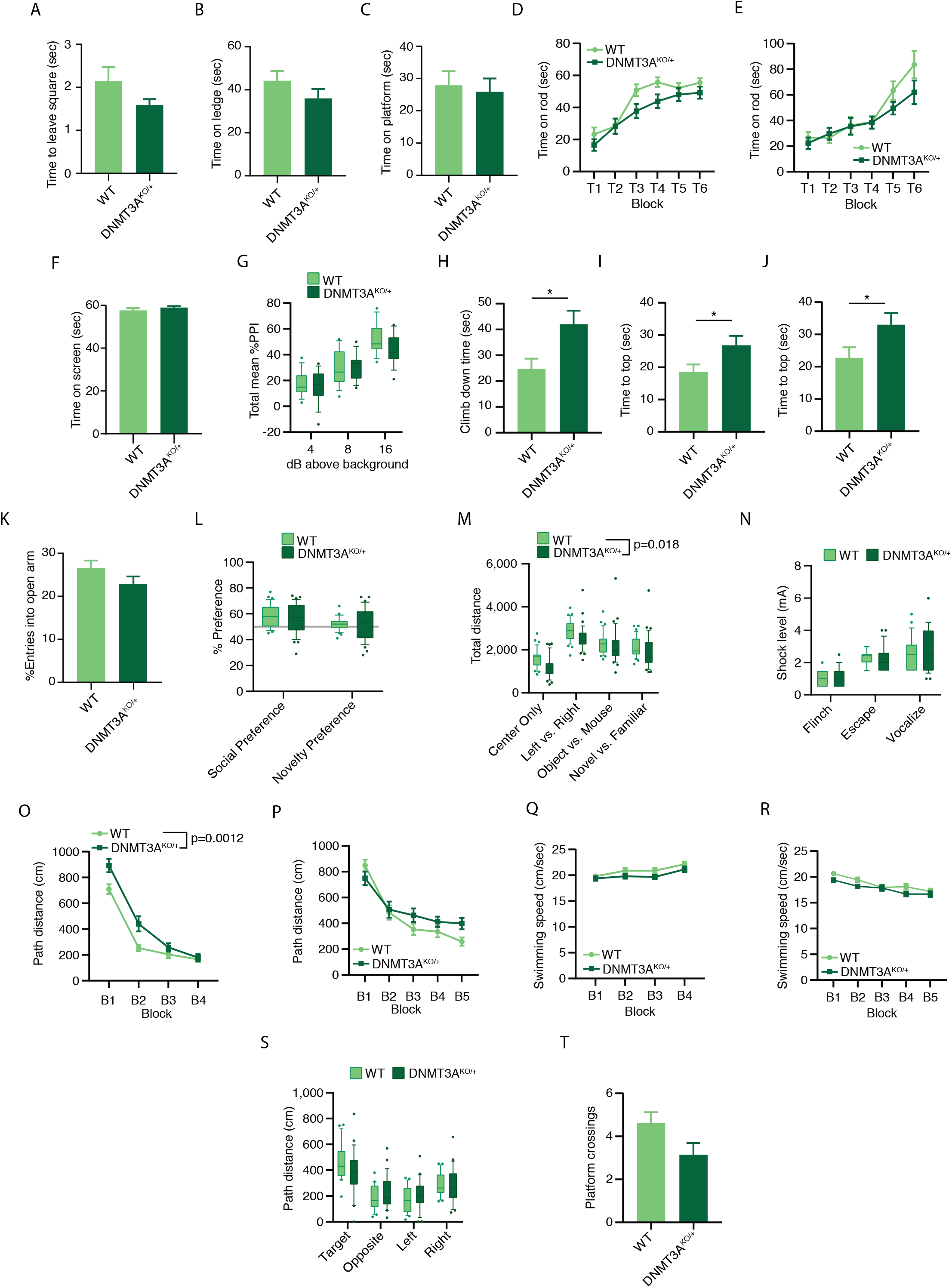
Related to Figure 3. (**A-J**) Comparison of DNMT3A^KO/+^ and WT mice across a battery of sensorimotor assays. DNMT3A^KO/+^ mice show no significant difference in (**A**) walking initiation, or latency to fall off (**B**) ledge or (**C**) platform. (**D-E**) DNMT3A^KO/+^ mice show no difference compared to WT littermates in motor coordination as evidenced by time on a continuous (**D**) and accelerating (**E**) rotarod. (**F**) DNMT3A^KO/+^ mice show no difference in grip strength compared to WT littermates evidenced by no change in time on an inverted screen. (**G**) Mean % pre-pulse inhibition shows no significant difference between genotypes. DNMT3A^KO/+^ mice show a significant increase in time to (**H**) climb down a pole (*P*=0.016, n=21,27; unpaired Student’s T-Test), and to the top of a (**I**) 60° inclined screen (*P*=0.039, n=21,27; unpaired Student’s T-Test) or a (**J**) 90° inclined screen (*P*=0.045, n=21,27; unpaired Student’s T-Test). (**K**) DNMT3A^KO/+^ mice show no deficit in elevated plus maze exploration as measured by percent entries into open arms. (**L**) Percent preference for novel conspecific in the 3-chambered social approach task for WT and DNMT3A^KO/+^ mice using time spent in zones as calculated: Mouse/(Mouse+Object)x100 or Novel/(Novel+Familiar)x100 within each animal. (**M**) Total distance traveled during the 3-chambered social approach task for WT and DNMT3A^KO/+^ mice shows a broad reduction of distance traveled across all trials by DNMT3A^KO/+^ mice (*P*=0.018 effect by genotype, F_(1,70)_=5.862, n=33,39; two-way ANOVA). (**N**) Shock sensitivity during conditioned fear test as indicated by the minimum shock needed to exhibit a behavioral response in mice shows no significant difference between genotypes. (**O-R**) Path distance to escape platform and swim speeds in the Morris water maze task. DNMT3A^KO/+^ mice show increased path distance to escape platform in both (**O**) cued trials (*P*=0.0012 effect by genotype, F_(1,46)_=11.93; *P*=0.0433 interaction effect of genotype and trial block, F_(3,138)_=2.784; n=21,27; two-way repeated-measures ANOVA) and (**P**) place trials (*P*=0.0408 interaction effect of genotype and trial block, F_(4,184)_=2.55 n=21,27; two-way repeated-measures ANOVA). No significant difference is seen in swimming speed during (**Q**) cued trials (*P=*0.0634 effect by genotype, F_(1,46)_=3.619 n=21,27; two-way repeated-measures ANOVA) and (**R**) place trials (*P=*0.098 effect by genotype, F_(1,46)_=2.845, n=21,27; two-way repeated-measures ANOVA). (**S**) DNMT3A^KO/+^ mice show no significant difference in time spent in the target quadrant of a Morris water maze compared to WT littermates. (**T**) DNMT3A^KO/+^ mice show a trend towards a reduction in platform crossings in the probe trial (*P*=0.0609; unpaired Student’s T-Test). Bar graphs and line plots indicate mean with SEM error bars. Box plots contain 10^th^-90^th^ percentiles of data, with remaining data represented as individual points.

**Figure S5.**
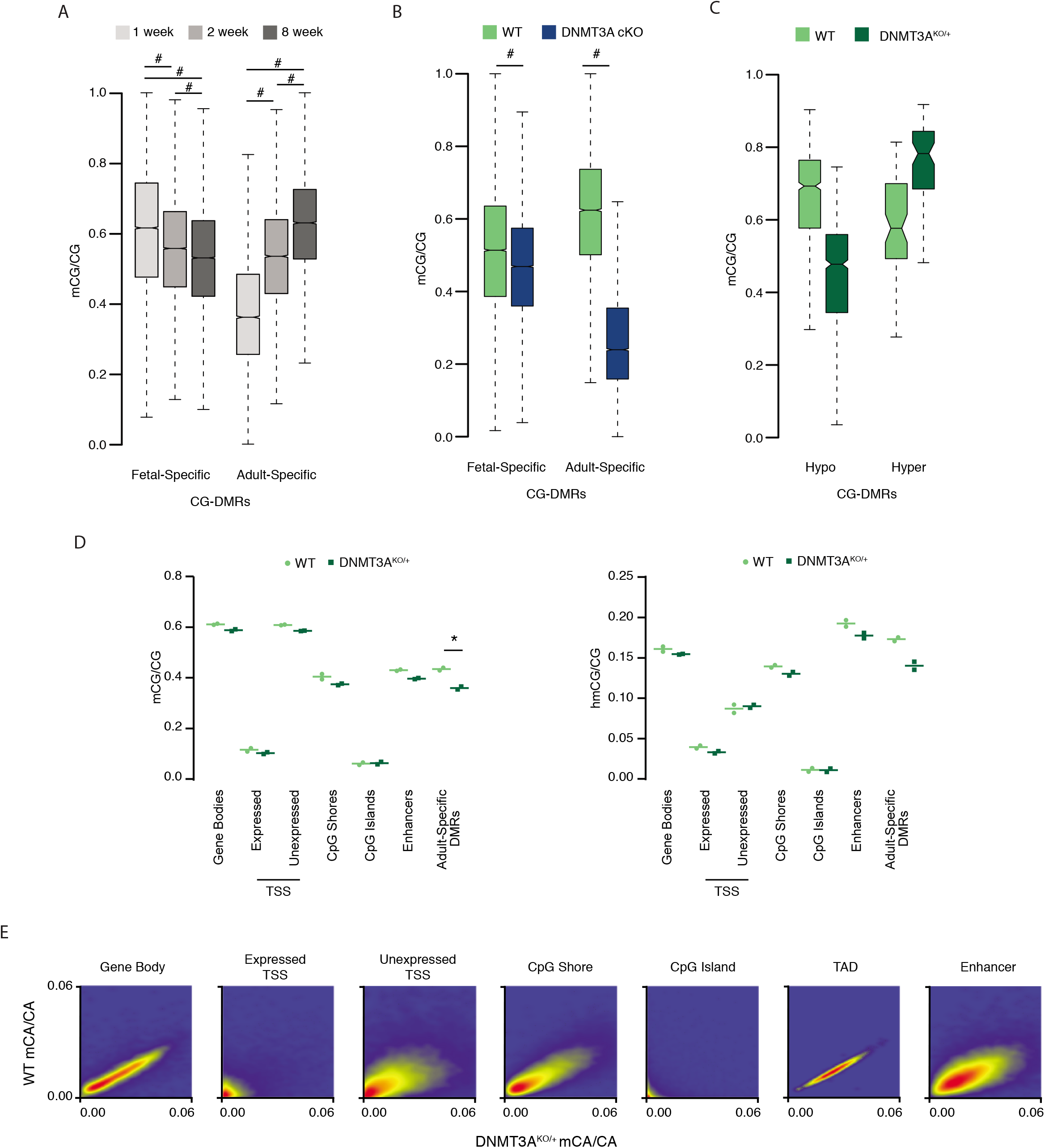
Related to Figures 4, 5, and 6. (**A**) Boxplots of CG methylation over postnatal development at regions called as having higher methylation in the frontal cortex of fetal versus adult tissue (Fetal-Specific DMR), or called as having higher methylation in the frontal cortex of the adult versus fetal tissue (Adult-specific DMR). DMRs from Lister et al. 2013, methylation data from Stroud et al. 2017. (#, *P*<2.2e-16; Wilcoxon rank sum test with Bonferroni correction) (**B**) Boxplots of cortical methylation of WT and DNMT3A cKO at 8 weeks postnatal within developmental DMRs shown in **A** (#, *P*<2.2e-16; Wilcoxon rank sum test with Bonferroni correction). (**C**) Boxplots of cortical methylation of WT and DNMT3A^KO/+^ at 8 weeks postnatally within DMRs defined in the DNMT3A^KO/+^ model. DMRs are called on this data set, so no additional statistics were run on genotype differences. (**D**) mCG/CG (left) and hmCG/CG (right) from WT and DNMT3A^KO/+^ cortices in various genomic contexts as measured by oxidative bisulfite sequencing. CpG islands were obtained from the UCSC table browser (Haeussler et al., 2019), and CpG Shores were defined as the 8kb surrounding them (*, *P*<0.05; paired Student’s T-Test with Bonferroni correction). (**E**) Smooth scatter plots of WT and DNMT3A^KO/+^ mCA/CA for classes of genomic regions. Box plots indicate median and quartiles.

**Figure S6.**
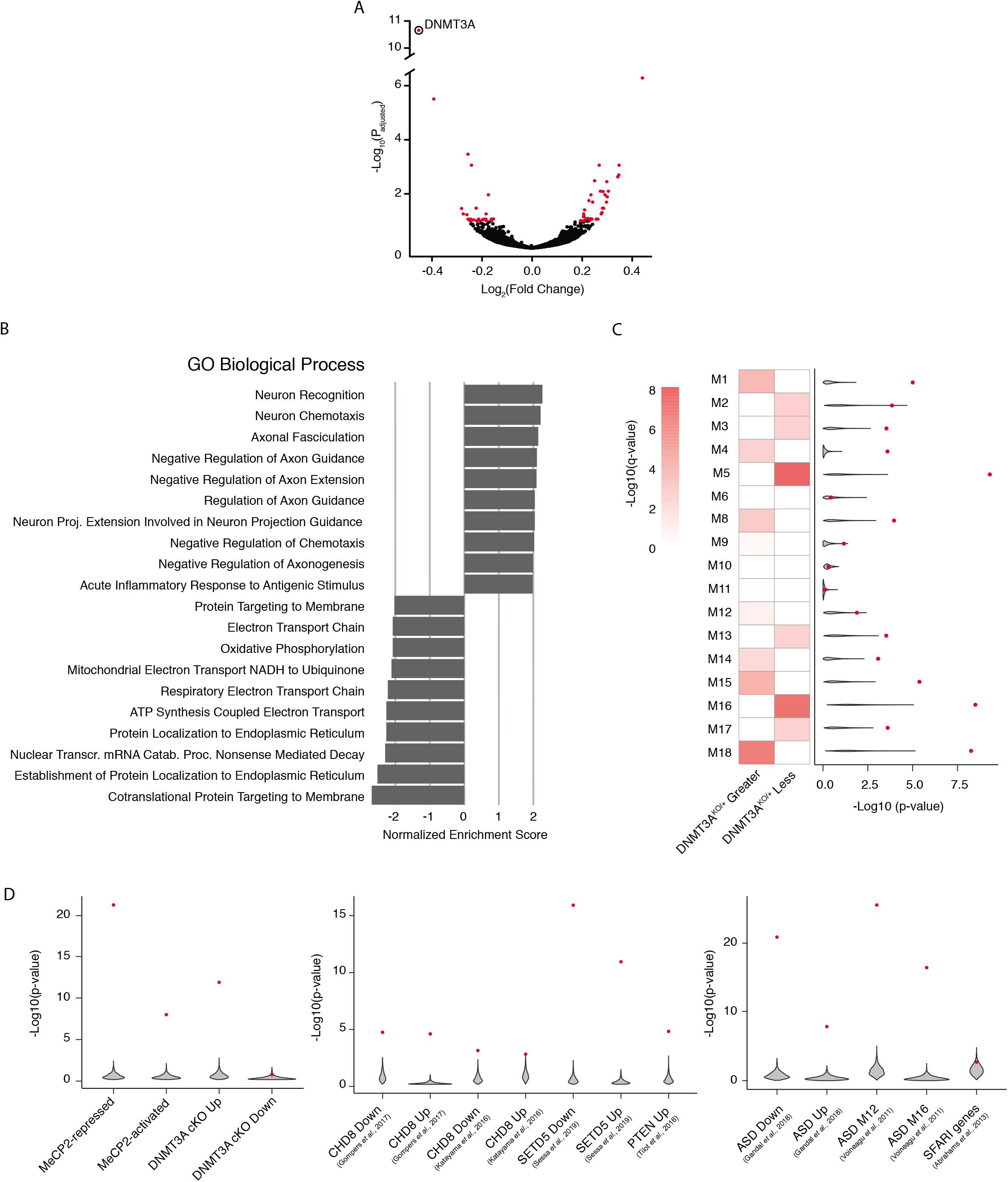
Related to Figures 6 and 7. (**A**) Volcano plot of DESeq log_2_ fold changes of the DNMT3A^KO/+^ versus WT. Genes reaching a significance of *p_adj_*.<0.1 are colored in red. (**B**) Top ten up- and down-regulated Gene Ontology terms from Broad GSEA Molecular Signatures Database version 7.0 (Subramanian et al., 2005). All terms are significant at an FDR<0.1. (**C**) GAGE analysis of developmental expression modules (Parikshak et al., 2013). Significant modules (q-value<0.1) are colored in red (left). Expression matched resampling of each gene set was performed 1,000 times and analyzed using GAGE for enrichment in DNMT3A^KO/+^ fold-change data (gray violin). This was compared with the true gene set p-value (red point) to test for significance (right). Only the direction of dysregulation in which the gene sets showed significance (i.e. DNMT3A^KO/+^ greater or less) is shown. (**D**) Expression matched resampling of GAGE analysis for gene sets displayed in Figures 6 and 7. Only the direction of dysregulation in which the gene set showed significance (i.e. DNMT3A^KO/+^ greater or less) is shown.

## STAR Methods

### Key Resources Table

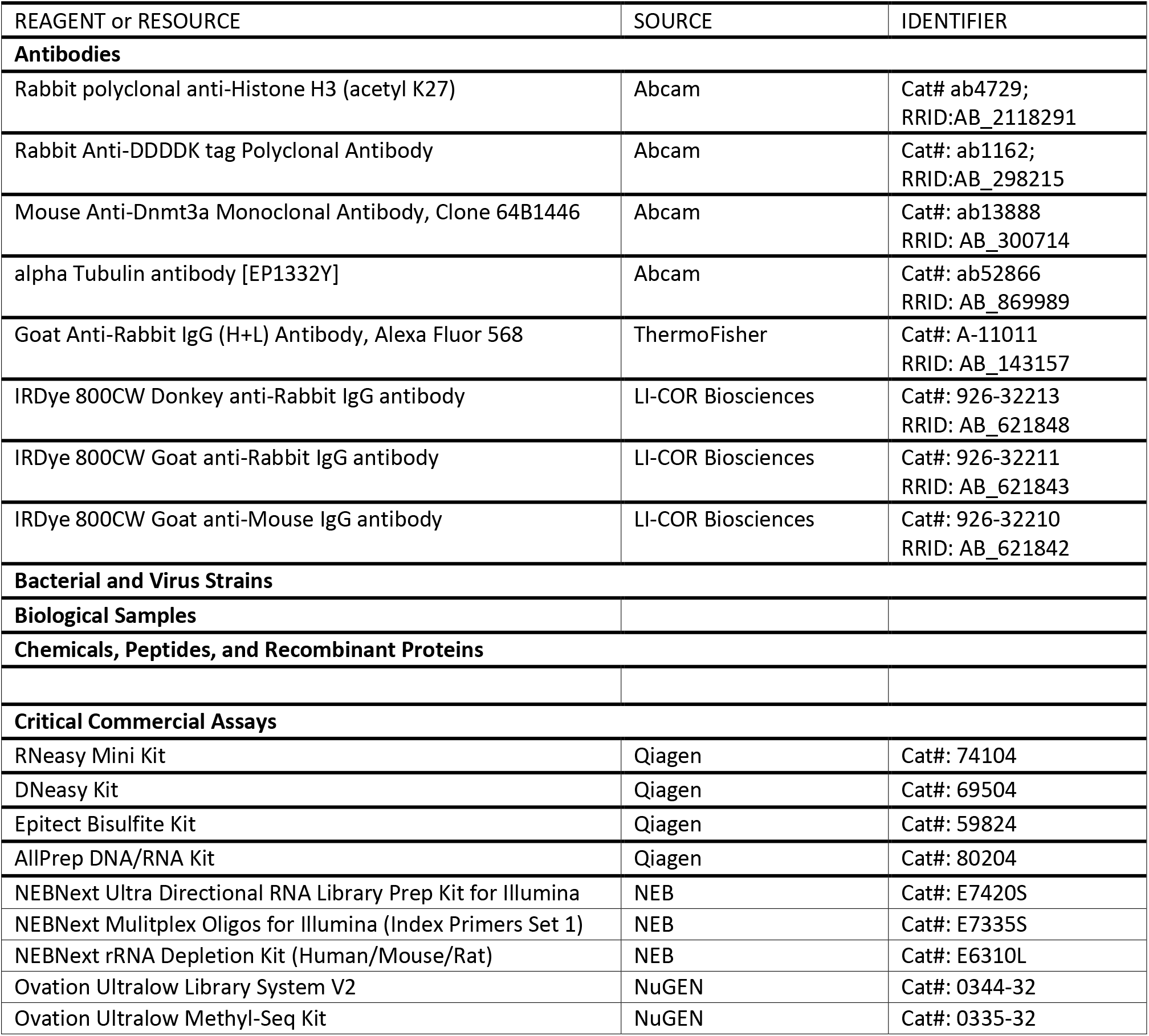
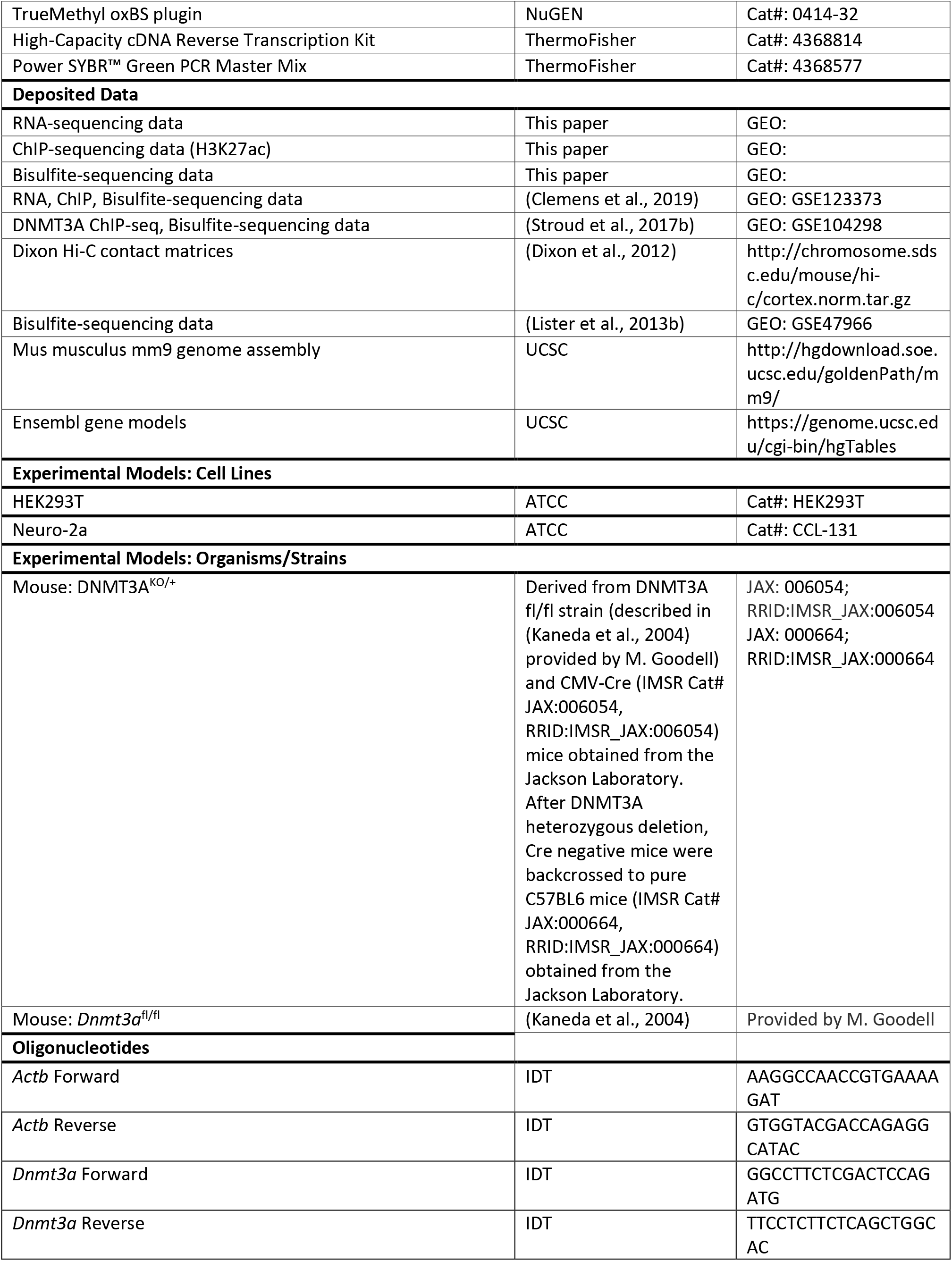
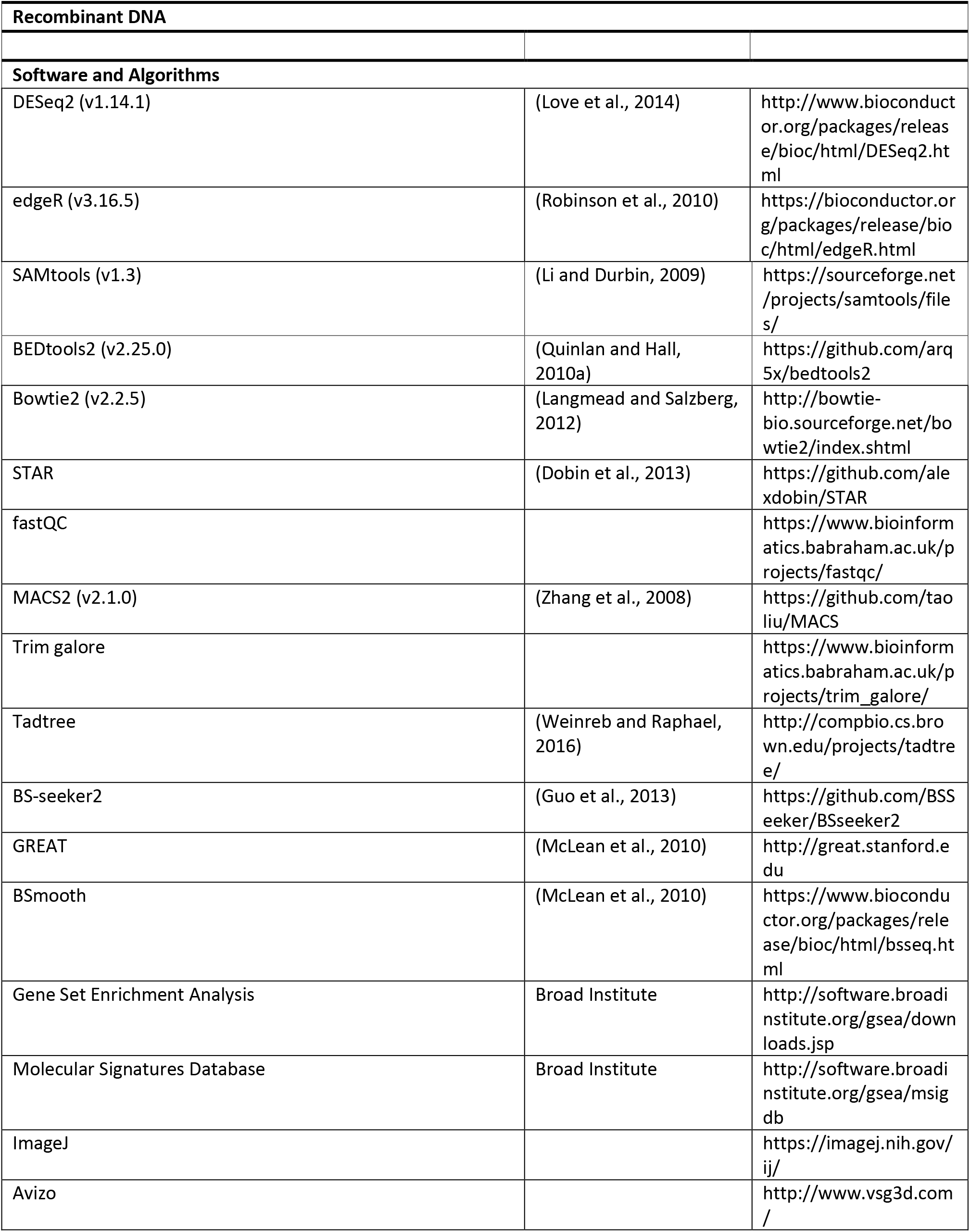
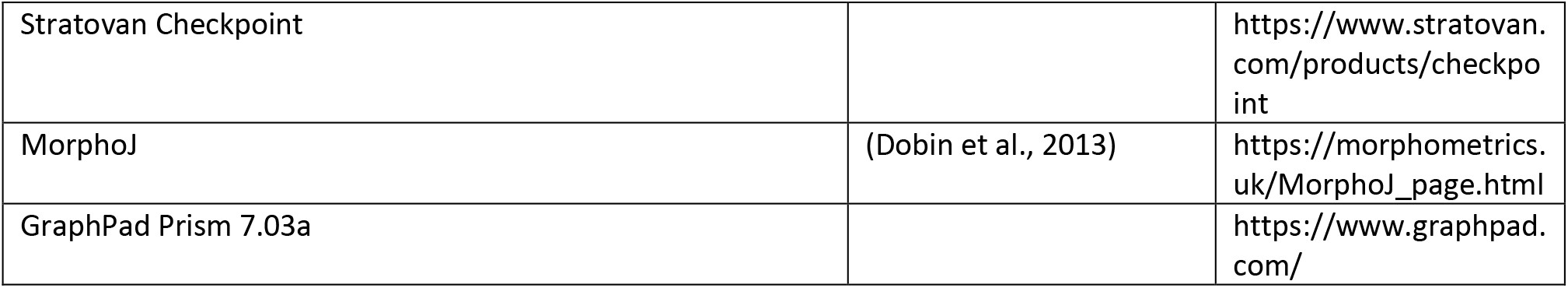

### Contact for Reagent and Resource Sharing

Requests for reagents and resources should be directed towards the Lead Contact, Harrison Gabel. This study did not generate new unique reagents.

### Experimental Model and Subject Details

#### Animal Husbandry

All animal protocols were approved by the Institutional Animal Care and Use Committee and the Animal Studies Committee of Washington University in St. Louis, and in accordance with guidelines from the National Institutes of Health (NIH). Mice were housed in a room on a 12:12 hour light/dark cycle, with controlled room temperature (20-22°C) and relative humidity (50%). Home cages measured 28.5 cm x 17.5 cm x 12 cm and were supplied with corncob bedding and standard laboratory chow and water. All mice were group-housed and adequate measures were taken to minimize animal pain or discomfort.

#### Transgenic animals

Male and female homozygous *Dnmt3a*^flx/flx^ mice (Kaneda et al., 2004) were bred together for viral-mediated DNMT3A replacement assay culture experiments. To generate the DNMT3A heterozygous mouse model, *Dnmt3a*^flx/flx^ mice were crossed to CMV:Cre (B6.C-Tg(CMV-cre)1Cgn/J) to generate *Dnmt3a^KO/+^Cre:CMV^+/-^* offspring. *Dnmt3a*^KO/+^Cre:CMV^+/-^ progeny were bred to C57BL/6J to outcross the cre recombinase and generate experimental genotype (DNMT3A^KO/+^). Subsequent experimental animals were generated from *Dnmt3a*^KO/+^ males mated to C57BL/6J females to generate *Dnmt3a*^KO/+^ and *Dnmt3a*^+/+^ experimental and control animals for experiments. *Dnmt3a*^KO/+^ females were not used for breeding to avoid social differences in mothering from mutant dams. Mice were genotyped with ear-DNA by PCR for *Dnmt3a* and *Cre*, and recombination was tested. Mice were weighed at a variety of timepoints to assess growth.

### Method Details

#### Immunocytochemistry

##### Staining

Neuro-2a cells (ATCC, CCL-131) were grown on coverslips and transfected with FLAG-tagged WT or mutant mouse DNMT3A plasmids and GFP plasmid. Coverslips were fixed with 4% paraformaldehyde in PBS for 20 minutes at room temperature, permeabilized with 0.1% Triton X-100 in PBS for 10 minutes at room temperature, and blocked with 1% BSA in PBS for 1 hour at room temperature. Coverslips were incubated overnight at 4°C in anti-DDDDK tag (FLAG-tag) primary antibody (Abcam, 1:5000, ab1162). Coverslips were then washed in PBS and incubated for 1 hour at room temperature with fluorescent secondary antibody (ThermoFisher, 1:500, A-11011) and counterstained with DAPI. *Imaging*. Images were captured using a Nikon A1Rsi confocal microscope with a 20x air objective. Laser settings were kept constant for each image. *Analysis/Quantification*. Cells were counted using an automatic threshold in FIJI and manually classified as displaying nuclear or non-nuclear signal by a blinded observer. This was determined by evaluating the overlap of FLAG signal (DNMT3A) with DAPI signal (nucleus). For mutants that did not reach expression levels comparable to the WT or for images that had too few positive cells, cell number was counted manually. 8 separate transfections were run, with each mutant being counted over 3 or more independent experiments. Sample sizes are as follows: WT, 15 images, 880 cells; W297del, 8 images, 435 cells; I310N, 8 images, 492 cells; S312fs11x, 12 images, 321 cells; G532S, 9 images, 695 cells; M548K, 7 images, 333 cells; V665L, 6 images, 635 cells; Y735C, 16 images, 613 cells; R749C, 8 images, 667 cells. P904L, 7 images, 692 cells. Percent nuclear was assessed per image and a generalized linear model was run comparing each mutant to WT. *P* values for each mutant were then Bonferroni corrected. We chose to use a generalized linear model with Bonferroni correction to allow for us to compare ratios of percent nuclear signal while taking into account experimental and biological replicates.

#### Modeling of DNMT3A disease mutations

HEK293T (ATCC, ACS-4500) or Neuro-2a cells (ATCC, CCL-131) were transfected with GFP and FLAG-tagged WT or mutant mouse DNMT3A plasmids. Collected cell lysates were ruptured by 3 freeze/thaw cycles using liquid nitrogen, or sonication ~42 hours after transfection. Samples were then either used for western blotting, the *in vitro* radioactive methyltransferase assay, or RNA isolation for qRT-PCR.

#### qRT-PCR

RNA was reverse transcribed using the High-Capacity cDNA Reverse Transcription Kit (Applied Biosystems). *Dnmt3a* and *Actb* were measured by qPCR using the Power SYBR™ Green PCR Master Mix and primers *Actb* (F:AAGGCCAACCGTGAAAAGAT, R:GTGGTACGACCAGAGGCATAC) and *Dnmt3a* (F:GGCCTTCTCGACTCCAGATG, R:TTCCTCTTCTCAGCTGGCAC). Relative quantity of *Actb* and *Dnmt3a* cDNA was determined by comparing the Ct of each primer set in each sample to a standard curve and then normalizing the DNMT3A signal by the ACTB signal. We chose to compare experimental conditions to WT samples using Student’s T-Tests with Bonferroni correction, as these are two normally distributed groups (visually checked) with similar variability.

#### *In vitro* radioactive methyltransferase assay

30 μl of cell lysate was used in the methyltransferase reaction previously described (Russler-Germain et al., 2014). Lysates were incubated at 37°C for 20 hours in 5 μl reaction buffer of 20 mM HEPES, 30 mM NaCl, 0.5 mM DTT, 1 mM EDTA, 0.2 mg/ml BSA, 5 mM 3 H-labeled SAM (PerkinElmer, NET155050UC) and 500 ng/μl Poly(dI-dC) substrate (Sigma P4929). Substrate was purified (Macherey-Nagel NucleoSpin Gel and PCR Clean-up) and radioactivity measured using a scintillation counter. In instances where DNMT3A mutant showed altered protein expression, cell lysate was re-balanced to match protein expression of WT DNMT3A. Only experimental replicates where WT DNMT3A showed a 1.5-fold increase compared to GFP alone were used for subsequent analysis. Outliers beyond 2 standard deviations above or below the mean were removed. Number of independent replicates are as follows: W297del, 18; I310N, 19; S312fs11x, 4; G532S, 10; M548K, 15; V665L, 11; Y375C, 13; R749C, 7; P904L, 14. Significance was assessed using a one-sample student’s t-test, as we are comparing groups normalized to WT and GFP back to the normalized value of 1.

#### Viral-mediated DNMT3A replacement assay

Functional activity of DNMT3A mutants in cortical neurons was determined by measuring methylation build-up *in vitro*. Timed mating of DNMT3A^flx/flx^ females and DNMT3A^flx/flx^ males was performed to collect embryonic cortical DNMT3A^flx/flx^ neurons at embryonic day 14.5. At E14.5, DNMT3A^flx/flx^ cortical neurons were isolated and plated (DIV 0). On DIV 3, neurons were either not perturbed or virally transduced with one of three conditions: 1) Cre only, 2) Cre and WT DNMT3A, or 3) Cre and mutant DNMT3A. DNA and RNA were isolated on DIV 12.5 using the AllPrep DNA/RNA Kit (Qiagen, 80204). DNA was used for whole genome bisulfite sequencing, and RNA was used for qRT-PCR for DNMT3A. Number of independent replicates are as follows: W297del, 5; S312fs11x, 4; G532S, 7; M548K, 6; V665L, 10; Y375C, 11; R749C, 6; P904L, 8. Buildup of methylation over development was done using timed mating of C57BL/6J mice and collected and prepared as described above. Significance was assessed using a one-sample student’s t-test, as we are comparing groups normalized to WT and GFP back to the normalized value of 1.

#### Ultrasonic vocalization and analysis

A total of 76 DNMT3A^KO/+^ (n=30, 16 male and 14 female) and litter-matched WT (n=46, 25 male and 21 female) mice were used for ultrasonic pup vocalization (USV) recording and analysis as previously described (Barnes et al., 2017; Dougherty et al., 2013; Holy and Guo, 2005). Dams were removed from the nest for a 10-minute acclimation, and individual pups had their body temperature measured using an infrared laser thermometer. Pups were then removed from their nest and placed in a dark, enclosed chamber. Ultrasonic vocalizations were recorded for 3 minutes with an Avisoft UltraSoundGate CM16 microphone and 416H amplifier using Avisoft Recorder software (gain = 6 dB, 16 bits, sampling rate = 250 kHz). Following this, pups were weighed and returned to their nest and littermates. All mice were recorded at postnatal days 5, 7, and 9, and on either day 11 or 15. Because not all animals were recorded from on day 11 or 15, only days 5, 7, and 9 were used for repeated measures ANOVA analysis. Frequency sonograms were prepared and analyzed in MATLAB (frequency range = 40 kHz to 120 kHz, FFT size = 256, overlap = 50%) with individual syllables identified and counted according to previously published methods (Dougherty et al., 2013; Holy and Guo, 2005). Significance was assessed using a within-subjects repeated measures ANOVA over timepoints 5-9, as these were when there was data from all experimental subjects, and these are optimal testing times where number of calls was highest.

#### Marble burying

A total of 27 DNMT3A^KO/+^ (n=13, 8 male and 5 female) and litter-matched WT (n=14, 7 male and 7 female) mice were used for marble burying. Marble burying is a natural murine behavior and has been used to indicate repetitive digging as well as anxiety-related behaviors. Protocol was adapted from previously published methods (Lazic, 2015; Maloney et al., 2019a). In brief, 8-week old mice were placed in a transparent enclosure (28.5 cm x 17.5 cm x 12 cm) with clean aspen bedding and 20 dark blue marbles evenly spaced in a 4 x 5 grid on top of the bedding. Animals were allowed to explore freely for 30 minutes. The number of buried marbles were counted every 5 minutes by two independent blinded observers. Marbles were considered “buried” if they were at least two-thirds covered by bedding. Enclosure and marbles were cleaned thoroughly between animals. Significance was assessed using a within-subjects repeated measures ANOVA to determine if rate of burying marbles is different between genotypes. These statistical methods are more appropriate than a simple t-test at 30 minutes, as mice may have buried all marbles before this timepoint, and significant changes in marble burying behavior may have occurred at an earlier timepoint in the assay.

#### Adult behavioral battery

A total of 72 DNMT3A^KO/+^ (n=39, 18 male and 21 female) and litter-matched WT (n=33, 15 male and 18 female) mice were used for adult behavioral testing. Mice were housed in mixed genotype home cages with 2-5 animals per cage, and all tests were performed during the light cycle. All experimenters were blinded to genotype during testing. For increased experimental rigor and reproducibility, we used three separate cohorts of mice to ensure quality and consistency in any observed phenotypes.

Testing started when mice were 3-4 months of age. The sequence of behavioral testing was designed to minimize carry-over effects across behavioral tests. Cohorts, ages, and testing order are in Supplementary Table 4. The majority of the assays were performed on cohorts 1 and 2 with cohort 3 being performed to test for reproducibility in some assays (Supplementary Table 4). Because of differences in testing sequences and exposure of mice to prior tests between cohorts, we examined separate cohorts individually and looked at combined cohorts (Supplementary Table 4). Testing was performed by the Washington University in St. Louis Animal Behavior Core.

#### One-hour locomotor activity

Locomotor activity was evaluated by computerized photobeam instrumentation in transparent polystyrene enclosures (47.6 cm x 25.4 cm x 20.6 cm) as previously described (Wozniak et al., 2004). Activity variables such as ambulations and vertical rearings were measured in addition to time spent in a 33 cm x 11 cm central zone.

#### Sensorimotor battery

Mice were assayed in walking initiation, balance (ledge and platform tests), volitional movement (pole and inclined screens), and strength (inverted screen) as previously described (Grady et al., 2006; Wozniak et al., 2004). For the walking initiation test, mice were placed on the surface in the center of a 21 cm x 21 cm square marked with tape and the time for the mouse to leave the square was recorded. During the balance tests, the time the mouse remained on an elevated plexiglass ledge (0.75 cm wide) or small circular wooden platform (3.0 cm in diameter) was recorded. During the Pole test, mice were placed at the top of a vertical pole with nose pointing upwards. The time for the mouse to turn and climb down the pole was recorded. For the inclined screen tests, a mouse was placed (oriented head-down) in the middle of an elevated mesh grid measuring 16 squares per 10 cm angled at 60° or 90°. Time for the mouse to turn 180° and climb to the top was recorded. For the inverted screen test, a mouse was placed on a similar screen and when the mouse appeared to have a secure grasp of the screen, the screen was inverted 180° and the latency for the mouse to fall was recorded. All tests had a duration of 60 seconds, except for the pole test which was 120 seconds. Two separate trials were done on subsequent days and averaged time of both trials was used for analysis. Data from the walking initiation, ledge, and platform tests were not normally distributed and therefore analyzed using Mann-Whitney U tests.

#### Continuous and accelerating rotarod

Motor coordination and balance were assessed using the rotarod test (Rotamex-5, Columbus Instruments, Columbus, OH) with three conditions: a stationary rod (60-second maximum), a rotating rod at constant 5 rpm (60-second maximum), and a rod with accelerating rotational speed (5 – 20 rpm, 180-second maximum) as previously described (Grady et al., 2006). This protocol is designed to minimize learning and instead measure motor coordination, so testing sessions were separated by 4 days to allow for extinction. Testing included one trial on stationary rod, and two trials on both the constant-speed rotarod and accelerating rotarod. Later timepoints in the constant speed rotarod test failed tests of normality, as the majority of mice stayed on the rotating rod for all 60 seconds. However, data were analyzed with two-way repeated-measures ANOVA.

#### Morris water maze

Spatial learning was assessed as previously described (Wozniak et al., 2004). Cued trials (visible platform, variable location) and place trials (submerged, hidden platform, consistent location) were conducted in which escape path latency, length, and swimming speeds were recorded. Animal tracking was done using a computerized system (ANY-maze, Stoelting). During cued trials, animals underwent 4 trials per day over 2 consecutive days with the platform being moved to a different location for each trial with few distal spatial cues available. Each trial lasted no longer than 60 seconds, with a 30-minute interval between each trial. Performance was analyzed across four blocks of trials (2 trials/block). After a three-day rest period, animals were tested on place trials, in which mice were required to learn the single location of a submerged platform with several salient distal spatial cues. Place trials occurred over 5 consecutive days of training, with 2 blocks of 2 consecutive trials (60-second trial maximum, 30-second inter-trial-interval after the mouse has reached the platform) with each block separated by 2 hours. Mice were released into different quadrants over different trials. Place trials were averaged over each of the five consecutive days (4 trials/block). One hour after the final block, a probe trial occurred (60-second trial maximum) in which the platform is removed, and the mouse is released from the quadrant opposite where the platform had been located. The time spent in pool quadrants, and the number of crossings over the exact platform location were recorded. DNMT3A^KO/+^ mice showed a small, but significant reduction in target zone time in cohort 2, though there was no difference in cohort 1 (Supplementary Table 4). Additionally, female mice had significantly faster swimming speeds than male mice across both genotypes (Supplementary Table 4). These findings and the observation that the DNMT3A^KO/+^ were slower moving on some of the sensorimotor tests, make path length a more appropriate variable than escape latency for evaluating performance, as escape latency can be confounded by the differences in swimming speeds.

#### 3-Chamber social approach

Sociability was assayed as previously described (Moy et al., 2004; Silverman et al., 2011). Mice were tested in a rectangular all-Plexiglas apparatus (each chamber measuring 19.5 cm x 39 cm x 22cm) divided into three chambers with walls containing rectangular openings (5 cm x 8 cm) and sliding doors. The apparatus was in a room with indirect light and was cleaned between tests with Nolvasan solution. Stimulus mice were contained within a small stainless-steel withholding cage (10 cm height x 10 cm diameter; Galaxy Pencil/Utility Cup, Spectrum Diversified Designs), allowing minimal contact between mice without allowing fighting. Between tests, withholding cages were cleaned with 75% ethanol solution. A digital video camera recorded movement of the mouse within the apparatus and allowed for tracking with ANY-maze (Stoelting). Distance and time spent in each chamber and investigation zones surrounding the withholding cages were recorded. Zones were defined as 12 cm in diameter from the center of withholding cages.

The test sequence consisted of 4 consecutive 10-minute trials in which the test mouse is placed in the middle chamber and allowed to freely explore the environment. In the first trial, the mouse is placed in the middle chamber with the doors to other chambers shut. In the second trial, the mouse is placed in the middle chamber and can explore all three chambers of the task, allowing it to acclimate to the environment. Neither genotype tested showed a preference towards a side of the chamber during this habituation. For the third trial, a sex-matched novel conspecific was placed within a withholding cage with the other cage remaining empty. For the fourth trial, the same sex-matched conspecific was in one withholding cage, while a new unfamiliar sex-matched stimulus mouse was placed in the other withholding cage. The locations of stimuli mice were counterbalanced across groups for the third trial and randomized novel or familiar for the fourth trial.

#### Elevated plus maze

Anxiety-like behaviors were examined using the elevated plus maze as previously described (Boyle, 2006). The apparatus contains a central platform (5.5 cm x 5.5 cm) with two opposing open arms and two opposing closed arms (each 36 cm x 6.1 cm x 15 cm) constructed of black Plexiglas. Mouse position is measured using beam-breaks from pairs of photocells configured in a 16 x 16 matrix and outputs are recorded using an interface assembly (Kinder Scientific) and analyzed using software (MotoMonitor, Kinder Scientific) to determine time spent, distance traveled, and entries made into open arms, closed arms, and the center area. Test sessions were conducted in a dimly lit room with each session lasting 5 minutes and each mouse tested over 3 consecutive days. Data shown are from day 1. All mice showed a decrease in time, distance, and entries into open arms on days 2 and 3. There was no significant difference between genotypes in percent entries into open arms (Figure S4K; *P*=0.137; unpaired Student’s T-Test) or total entries into arms (data not shown), indicating that both genotypes explored the maze. Percent distance traveled in open arms showed similar effects to percent time in open arms (Percent distance traveled: *P*=0.027; unpaired Student’s T-Test). Analysis of these data in individual cohorts detected DNMT3A^KO/+^ significant effects for the percent of open arm time on the first day in cohorts 1 and 3, with no evidence of an effect in cohort 2 (Supplementary Table 4). Individual cohorts also showed no significant difference between genotypes in percent open arm entries (Supplementary Table 4) suggesting that mice explored the elevated plus maze sufficiently to detect anxiety-like behaviors.

#### Acoustic startle/prepulse inhibition

Sensorimotor gating was evaluated as previously described (Dougherty et al., 2013; Gallitano-Mendel et al., 2008; Hartman et al., 2001). In short, mice were presented with an acoustic startle response (120 dB auditory stimulus pulse, 40 ms broadband burst) and a pre-pulse (response to pre-pulse plus startle pulse). Stimulus onset began at 65 seconds, and 1ms force readings were obtained and averaged to produce an animal’s startle amplitude. 20 startle trials were presented in 20 minutes. The first 5 minutes were an acclimation period where no stimuli above the 65 dB background were presented. The session started and finished with 5 consecutive startle (120 dB pulse) trials. The middle 10 trials were interspersed with pre-pulse trials, consisting of an additional 30 presentations of 120 dB startle stimuli preceded by pre-pulse stimuli of 4, 12, or 20 dB above background (10 trials for each PPI trial type). To calculate percent pre-pulse inhibition, we used %PPI = 100 × (ASR_startle pulse alone_ – ASR_prepulse + startle pulse_)/ASR_startle pulse alone_.

#### Conditioned fear

Fear conditioning was done as previously described (Maloney et al., 2019a, 2019b). Mice were habituated to an acrylic chamber (26 cm x 18 cm x 18 cm) containing a metal grid floor and an odorant and was illuminated by LED light which remained on for the duration of the trial. Day 1 testing lasted 5 minutes in which an 80 dB tone sounded for 20 seconds at trial timepoints 100, 160, and 220 seconds. A 1.0 mA shock (unconditioned stimulus) occurred within the last 2 seconds of the tone (conditioned stimulus). Baseline freezing behavior during the first 2 minutes and the freezing behavior during the last 3 minutes was quantified using image analysis (Actimetrics, Evanston, Illinois). On Day 2, testing lasted for 8 minutes in which the light was illuminated but no tones or shocks were presented. On Day 3, testing lasted for 10 minutes in which the mouse was placed in an opaque chamber with a different odorant than the original test chamber. The 80 dB tone began at 120 seconds and lasted for the remainder of the trial and freezing behavior to the conditioned auditory stimulus was quantified for the remaining 8 minutes. Small elevated freezing levels of the DNMT3A^KO/+^ mice for the contextual fear and auditory cue data could be interpreted as evidence for an increased baseline propensity to freeze or stronger fear conditioning in the mutant mice. However, an alternative hypothesis is that the exaggerated freezing levels displayed by the DNMT3A^KO/+^ mice may reflect an emotional hypersensitivity to the footshock as was documented by their freezing levels during tone-shock training. In support of the latter hypothesis, evaluation of baseline freezing levels in individual cohorts showed that they were only significantly different in one of the two cohorts tested.

#### Statistical analysis for behavioral tests

Behavioral data were analyzed with R v3.3.2 (including the ANOVA function from the Car package in R (Fox and Weisberg, 2011)) and plots were made using GraphPad Prism 7.03a. Normality was assessed using the Shapiro-Wilkes test and visually confirmed. Data not normally distributed were analyzed using non-parametric tests, with the exception of continuous rotarod data. Sexes were considered separately with genotype and times/block as fixed factors with no consistent sex effects observed, therefore data were collapsed across sex. Statistical testing was performed using planned assay-specific methods, such as using Student’s T-Tests for single parameter comparisons between genotypes, and within-subjects two-way repeated-measures ANOVA for comparisons across timepoints. Individual timepoints within repeated measures tests were evaluated using Sidak’s multiple comparisons test. Individual cohorts were analyzed separately and in aggregate with similar trends seen across cohorts (Supplementary Table 4), therefore data from all cohorts were included together.

#### Tissue

Brain tissue was dissected from DNMT3A^KO/+^ and WT littermate mice in ice-cold PBS, flash-frozen in liquid nitrogen, and stored at −80°C.

#### Western blotting

##### Western blotting from cell culture

Neuro-2a or HEK293T cells were collected and combined with 2x laemmli buffer with 5% β-mercaptoethanol. Samples were passed through a Wizard Column (Fisher, Wizard Minipreps Mini Columns, PRA7211), boiled for 5 minutes, and run on a BioRad 4-12% acrylamide gel at 125 V for 60 minutes. Samples were then transferred to a nitrocellulose membrane, which was bisected between 37kDa and 50kDa bands. Membranes were blocked with 3% bovine serum albumin in TBS-T for 1 hour at room temperature and then the lower membrane was immunostained with anti-GFP (ThermoFisher, 1:2000, A-11122) while the upper membrane was immunostained with anti-DDDDK (Abcam, 1:1000, ab1162) for 12-16 hours at 4°C. All primary and secondary antibodies were diluted in 3% Bovine Serum Albumin in TBS-T. Membranes were then washed with TBS-T then incubated for 1 hour at room temperature with IR-dye secondary antibody (IRDye 800CW Donkey anti-Rabbit, LI-COR Biosciences, 1:15,000, product number: 926-32213). Blots were then washed in PBS, and imaged using the LiCOR Odyssey XCL system, and quantified using Image Studio Lite software (LI-COR Biosciences). FLAG (DDDDK) and GFP levels were normalized to a standard curve, and protein levels are expressed as normalized DDDDK values divided by normalized GFP values to enable comparison of FLAG (DDDDK) levels between blots. Each blot included a standard curve and WT samples. Outliers beyond 2 standard deviations above or below the mean were removed. Number of independent replicates are as follows: WT, 29; W297del, 7; I310N, 7; S312fs11x, 12; G532S, 7; M548K, 9; V665L, 7; Y375C, 8; R749C, 6; P904L, 7. Significance was assessed using a one sample T-Test, as protein expression levels were normalized to GFP and WT, and mutant protein expression was compared to the normalized WT value of 1.

##### Western blotting from tissue

Brain tissue samples were homogenized with a dounce homogenizer in buffer with protease inhibitors (10mM HEPES pH 7.9, 10mM KCl, 1.5mM MgCl2, 1mM DTT, 10mM EDTA). A portion of the lysate was removed and 1% SDS was added. Samples were boiled for 10 minutes, followed by a 10-minute spin at 15,000g. Supernatant was collected and run through a Wizard Column (Fisher, Wizard Minipreps Mini Columns, PRA7211), then diluted in LDS sample buffer with 5% β-mercaptoethanol. Samples were boiled for 5 minutes, run on an 8% acrylamide gel for 60 minutes at 125 V, and transferred to a nitrocellulose membrane. Membrane was bisected between 75kDa and 100kDa. Membranes were blocked with 3% bovine serum albumin in TBS-T for 1 hour at room temperature, and the upper membrane was immunostained with anti-DNMT3A (Abcam, 1:1000, ab13888) while the lower membrane was immunostained with anti-α-Tubulin (Abcam, 1:1000, ab52866) for 12-16 hours at 4°C. All primary and secondary antibodies were diluted in 3% Bovine Serum Albumin in TBS-T. Membranes were then washed with TBS-T then incubated for 1 hour at room temperature with IR-dye secondary antibody (IRDye 800CW Goat anti-Rabbit, or IRDye 800CW Goat anti-Mouse, LI-COR Biosciences, 1:15,000, product numbers: 926-32211 and 926-32210 respectively). Blots were then washed in PBS, imaged using the LiCOR Odyssey XCL system, and quantified using Image Studio Lite software (LI-COR Biosciences). DNMT3A and α-Tubulin levels were normalized to a standard curve, and protein levels are expressed as normalized DNMT3A values divided by normalized α-Tubulin values to enable comparison of DNMT3A levels between blots. For brain region analysis, sample sizes of n=4 per genotype (2 male and 2 female pairs) were used. For time course analysis, sample sizes of n=2 per genotype (1 male and 1 female pairs) were used for all time points except the 2-week timepoint in which n=6 (3 male and 3 female pairs) was used. Significance was assessed using a two way ANOVA considering genotype and time to determine if there was a detectable difference in protein expression over time.

#### Bisulfite sequencing

##### Whole genome bisulfite sequencing from cortical cultures

Samples were chosen for whole genome bisulfite sequencing if mutant and WT samples expressed equal amounts of *Dnmt3a* mRNA as measured by qRT-PCR (Figure S2D). DNA from cortical cultures was bisulfite converted and prepared for sequencing using the Tecan Ovation Ultralow Methyl-Seq Kit (Tecan, 0335-32) and the Epitect Bisulfite Kit (Qiagen, 59824) was used for bisulfite conversion. We used alternate bisulfite conversion cycling conditions ([95°C, 5 min; 60°C, 20 min] x 4 cycles, 20°C hold) to ensure lowest possible bisulfite non-conversion rate. Libraries were PCR-amplified for 10-11 cycles. Libraries were then pooled and sequenced at a depth of 0.01-0.03x genomic coverage using an Illumina MiSeq 2×150 through the Spike-In Cooperative at Washington University in St. Louis. Significance was assessed using a one-sample student’s t-test, as we are comparing groups normalized to WT and GFP back to the normalized value of 1.

##### Whole genome bisulfite sequencing from tissue

DNA was isolated from tissue using the DNEasy Kit (Qiagen). 300 ng of DNA was prepared for sequencing using the Ovation Ultralow Methyl-Seq Kit (Tecan, 0335-32) with and the Epitect Bisulfite Kit (Qiagen, 59824) was used for bisulfite conversion. For these samples, 300 ng of DNA was fragmented for 45 seconds with the Covaris E220 sonicator (10% Duty Factory, 175 Peak Incidence Power, 200 cycles per burst, milliTUBE 200μL AFA Fiber). DNA was then purified using 0.7 volumes of Agencourt Beads to select for long DNA inserts for sequencing. We used alternate bisulfite conversion cycling conditions ([95°C, 5 min; 60°C, 20 min] x 4 cycles, 20°C hold) to ensure lowest possible bisulfite non-conversion rate. Libraries were PCR-amplified for 12 cycles. Libraries were then pooled and sequenced using an Illumina MiSeq 2×150 through the Spike-In Cooperative at Washington University in St. Louis. Samples for shallow-depth sequencing (Figure 4A,B) were sequenced at 0.01-0.03x genomic coverage. For brain region and liver methylation, n=2 per genotype per region (one male pair, one female pair). For developmental time course methylation, n=3-4 per genotype per timepoint, with at least one male and one female pair. 8-week cortex samples for deep sequencing (Figure 5, Figure 6A,B,D, Figure S5C-E) were sequenced at 6.4-7.8x coverage per biological rep (two technical reps per biological rep, two biological reps per genotype) using only male genotype pairs. For shallow sequencing experiments, significance was assessed using either a two-sample Student’s T-Test to compare global methylation values of the cortex between two genotypes, or using a two way ANOVA to compare broad methylation changes across a variety of brain regions between genotypes. Genomic element comparisons were done using two-sample Student’s T-Tests with Bonferroni correction.

##### Oxidative bisulfite sequencing from tissue

DNA was isolated from tissue using the DNEasy Kit (Qiagen, 69504). 450 ng of DNA was prepared for sequencing using the Ovation Ultralow Methyl-Seq Kit (Tecan, 0335-32) with TrueMethyl oxBS plugin (Tecan, 0414-32). For these samples, 450 ng of DNA was fragmented for 45 seconds with the Covaris E220 sonicator (10% Duty Factory, 175 Peak Incidence Power, 200 cycles per burst, milliTUBE 200μL AFA Fiber). DNA was then purified using 0.7 volumes of Agencourt Beads to select for long DNA inserts for sequencing. 300 ng of DNA was used for OxBS libraries, whereas the remaining 150 ng of DNA was used for bisulfite libraries. We used alternate bisulfite conversion cycling conditions ([95°C, 5 min; 60°C, 20 min] x 2 cycles; 95°C, 5 min; 60°C, 40 min; 95°C, 5 min; 60°C, 45 min; 20°C hold) to ensure lowest possible bisulfite non-conversion rate. Bisulfite and oxidative bisulfite libraries were PCR-amplified for 11 and 13 cycles respectively. Libraries were then pooled and sequenced using an Illumina MiSeq 2×150 through the Spike-In Cooperative at Washington University in St. Louis. Samples were sequenced at 0.8-2.2x genomic coverage per replicate (two replicates per genotype). Genomic element comparisons were done using two-sample Student’s T-Tests with Bonferroni correction.

#### Whole-genome bisulfite analysis

Bisulfite sequencing analysis was performed as previously described (Clemens et al., 2019). Briefly, data were adapter-trimmed, mapped to mm9, then deduplicated and called for methylation using BS-seeker2. Methylation levels across regions were assessed using bedtools map -o sum, summing the number of reads mapping to Cs (supporting mC) and the amount of coverage in the region, then dividing those two numbers(Quinlan and Hall, 2010b). Hydroxymethylation was calculated as the percent methylation found in the BS-seq minus the percent methylation found in the matching oxBS-seq. Due to count noise, this occasionally resulted in apparent negative hydroxymethylation. During bisulfite sequencing not all DNA can be efficiently bisulfite converted. Though our methods should maximize the amount of converted unmethylated C, there is still a small percentage of unmethylated cytosines that are called as methylated due to non-conversion (0.2-0.3%). Due to this non-conversion, very lowly methylated regions (e.g. mCA at CpG islands) may not show the same percent reduction in mCA as highly methylated regions. Data were visualized using the UCSC genome browser (http://genome.ucsc.edu) (Kent et al., 2002). Average methylation per-sample is normally distributed in all regions examined, and variance between genotypes is similar, fitting the assumptions of a 2-sample t-test. Methylation levels for individual elements are not necessarily normally distributed, so non-parametric tests were used instead.

#### Differentially methylated region detection

BSmooth (Hansen et al., 2012) was used to call differentially CpG methylated regions between DNMT3A^KO/+^ and WT mice, using two technical replicates each of two biological replicates. CG sites were filtered for requiring at least 2x coverage in all replicates and differentially methylated regions were called with a statistical threshold of t-stat >2.0. These regions were further filtered for a length >100 bp and a requirement that the smoothed per-rep methylation values were consistent. For hypomethylated regions, all WT mCG/CG values needed to be greater than any KO mCG/CG value and all KO methylation values needed to be higher than all WT methylation values for hypermethylated regions. Data fit the assumptions and requirements of BSmooth. Data were distributed evenly between chromosomes, and the overlap between DMRs and regions of interest fit a hypergeometric distribution, making a fisher’s exact test appropriate.

#### RNA sequencing

Total RNA isolation was carried out as previously described (Clemens et al., 2019). In brief, cerebral cortex was dissected in ice-cold PBS from DNMT3A^KO/+^ and WT littermates at 8 weeks of age (n=7 pairs, 3 male, 4 female). Cortex was lysed in RLT buffer following the RNeasy Mini Kit (Qiagen, 74104). RNA libraries were generated from 250 ng of RNA with NEBNext Ultra Directional RNA Library Prep Kit for Illumina (NEB) using a modified amplification protocol (37°C, 15 minutes; 98°C, 30 seconds; [98°C, 10 seconds; 65°C, 30 seconds; 72°C, 30 seconds]x13; 72°C, 5 minutes; 4°C hold). RNA libraries were pooled at a final concentration of 10nM and sequenced using Illumina HiSeq3000 1×50bp with the Genome Technology Access Center at Washington University in St. Louis, typically yielding 15-30 million single-end reads per sample.

#### RNA sequencing analysis

RNA sequencing analysis was performed as previously described (Clemens et al., 2019). Briefly, raw FASTQ files were trimmed with Trim Galore and rRNA sequences were filtered out with Bowtie. Remaining reads were aligned to mm9 using STAR (Dobin et al., 2013) with the default parameters. Reads mapping to multiple regions in the genome were then filtered out, and uniquely mapping reads were converted to BED files and separated into intronic and exonic reads. Finally, reads were assigned to genes using bedtools coverage-counts (Quinlan and Hall, 2010b).

For gene annotation we defined a “flattened” list of longest transcript forms for each gene, generated on Ensgene annotations and obtained from the UCSC table browser. For each gene, Ensembl IDs were matched up to MGI gene names. Then, for each unique MGI gene name, the most upstream Ensgene TSS and the most downstream TES were taken as that gene’s start and stop. Based on these Ensembl gene models, we defined TSS regions and gene bodies. Differentially expressed genes were identified using a Wald test through DESeq2, running using default parameters on exonic reads from the DNMT3A^KO/+^ and WT.

#### Chromatin immunoprecipitation protocol

Chromatin immunoprecipitation was performed as previously described (Clemens et al., 2019; Cohen et al., 2011). Cerebral cortex was dissected on ice in PBS from DNMT3A^KO/+^ and WT littermates at 8-weeks old (n=5 pairs, 3 male, 2 female). The tissue was flash-frozen in liquid nitrogen and stored at −80°C. Chromatin were fragmented with the Covaris E220 sonicator (5% Duty Factory, 140 Peak Incidence Power, 200 cycles per burst, milliTUBE 1mL AFA Fiber). ChIP was performed with H3K27ac antibody (0.025-0.1μg; Abcam, ab4729) and libraries were generated using Ovation Ultralow Library System V2 (Tecan, 0344NB-32). Libraries were pooled to a final concentration of 8-10nM and sequenced using Illumina HiSeq 3000 with the Genome Technology Access Center at Washington University in St. Louis, typically yielding 15-40 million single-end reads per sample.

#### Chromatin immunoprecipitation analysis

ChIP sequencing analysis was performed as previously described (Clemens et al., 2019). Briefly, reads were mapped to mm9 using bowtie2 and reads were extended based on library sizes and deduplicated. Bedtools coverage–counts was used to quantify ChIP signal at the transcriptional start site (TSS), gene body (GB), and transcriptional end site (TES) (Quinlan and Hall, 2010b). edgeR was then used to determine differential ChIP-signal across genotypes. Data were visualized using the UCSC genome browser (http://genome.ucsc.edu) (Kent et al., 2002).

#### Controlled resampling

A similar resampling approach was used as previously described (Clemens et al., 2019). Briefly, for every entry in a sample set (e.g. DNMT3A-dysregulated genes), an entry in the control set (e.g. all other genes) with a similar desired characteristic (e.g. expression) was selected, generating a control set of the same size and variable distribution as the sample set.

#### Identification of dysregulated enhancers

Enhancer regions from Clemens et al. 2019 were used, and enhancers dysregulated in the DNMT3A^KO/+^ were called using the same method. Briefly, H3K27ac ChIP-seq reads were quantified in all acetyl peak regions, and edgeR was used to identify peaks with significantly different amounts of H3K27ac signal. Peak regions were then divided into promoters, enhancers, and non-identified peaks. Data fits the assumptions of BSmooth. Overlap between misregulated enhancers in different genotypes fit a hypergeometric distribution.

#### GAGE

Gene set enrichment analysis for the gene sets described was performed using the Generally Applicable Gene-set Enrichment (GAGE) program (Luo et al., 2009). Analysis was performed directionally on input of shrunken, log-normalized exonic fold changes output from DESeq2 analysis of DNMT3A^KO/+^ versus WT RNA-seq data. Gene sets with an FDR q-value below 0.1 and an adjusted p-value below 0.5 following expression matched resampling repeated 1,000 times were considered statistically significant. Gene sets were selected for analysis from both human and mouse studies of autism associated genes. SFARI genes (Abrahams et al., 2013) with scores of equal to or less than 3 were considered. Date accessed: 6/20/2019.

#### GSEA

Gene Set Enrichment Analysis (GSEA) (version 7.0, the Broad Institute of MIT and Harvard, http://software.broadinstitute.org/gsea/downloads.jsp) was performed on shrunken, log-normalized exonic fold changes from DESeq2 between DNMT3A^KO+^ and WT RNA-seq data. GSEA calculated a gene set Enrichment Score (ES) that analyzed genes were enriched in the biological signal conduction on the MsigDB (Molecular Signatures Database, http://software.broadinstitute.org/gsea/msigdb). Background was set to all expressed genes in this study and 1,000 permutations were set to generate a null distribution for enrichment score in the hallmark gene sets and functional annotation gene sets. The gene sets database used for enrichment analysis were ‘c5.all.v7.0.symbols.gmt’, ‘c5.bp.v7.0.symbols.gmt’, ‘c5.cc.v7.0.symbols.gmt’and ‘c5.mf.v7.0.symbols.gmt’ and FDR <0.1 was defined as the cut-off criteria for significance.

#### Craniofacial morphological analyses

A total of 24 sex-matched littermate paired mice (DNMT3A^KO/+^ n=12, 7 male, 5 female; WT n=12, 7 male, 5 female) across 3 time-points (8 weeks DNMT3A^KO/+^ n=4, WT n=4; 20 weeks DNMT3A^KO/+^ n=4, WT n=4; 25 weeks DNMT3A^KO/+^ n=4, WT n=4) were fixed in 4% paraformaldehyde through intracardiac perfusions. Whole mouse heads were scanned at the Musculoskeletal Research Center at Washington University in St. Louis using a Scanco μCT40 machine. CT images had voxel dimensions of 0.018 millimeters and were reconstructed on a 2048×2048 pixel grid. The CT images were converted to 8bit images using ImageJ (https://imagej.nih.gov/ij/) and surface reconstructions were acquired in Avizo (http://www.vsg3d.com/). Thirty-five three-dimensional landmarks were collected from surface reconstructions of the cranium and mandible using Stratovan Checkpoint (https://www.stratovan.com/products/checkpoint).

Generalized Procrustes Analysis in MorphoJ software was used to explore the differences and similarities of shape between the DNMT3A^KO/+^ mice and their WT littermates as previously described (Hill et al., 2013). To control for possible differences in size, the landmark coordinate data were natural log-transformed and analyzed with a linear regression model. Additionally, to localize differences in form to specific linear distances, landmark data were analyzed using Euclidean Distance Matrix Analysis (EDMA).

#### Bone length measurements

A total of 24 sex-matched littermate paired mice (DNMT3A^KO/+^ n=12, 7 male, 5 female; WT n=12, 7 male, 5 female) across 3 time-points (8 weeks DNMT3A^KO/+^ n=4, WT n=4; 20 weeks DNMT3A^KO/+^ n=4, WT n=4; 25 weeks DNMT3A^KO/+^ n=4, WT n=4) were fixed in 4% paraformaldehyde through intracardiac perfusions. Decapitated mouse bodies were scanned at the Musculoskeletal Research Center at Washington University in St. Louis using a Faxitron Model UltraFocus100 Dual X-Ray machine. Bone lengths were measured using ImageJ. Data were taken over three age time-points: 8 weeks, 20 weeks, and 25 weeks of age for male and female mice. There was no significant difference in bone lengths based upon sex, but there was a difference based by age. To normalize for this age effect, data were expressed as DNMT3A^KO/+^ bone lengths normalized to the WT lengths within groups. Left and right bones were measured and the larger was used for analysis.

#### Experimental design

Authenticated cell lines from ATCC (HEK293T, NEURO2A) were used, and no mycoplasma contamination testing was needed. Sample sizes were chosen based upon previously published studies using similar techniques. Statistical tests and exclusion criteria (values beyond 2 standard deviations of the group mean) were similar to that of previously published studies and indicated in the appropriate methods. For all animal experiments, experimenters were blinded to genotype during data collection. No treatment conditions were used, so no samples or animals were allocated to experimental groups and no randomization was needed. Tests that assume equal variance were only run if group variances were similar, otherwise alternative tests were used.

#### Data availability statement

The data that support the findings of this study are available from the corresponding author upon request. DOIs for all published gene sets used in comparison and enrichment analysis: Lister et al. 2013: https://doi.org/10.1126/science.1237905;

Clemens et al. 2019: https://doi.org/10.1016/j.molcel.2019.10.033;

Stroud et al. 2017: https://doi.org/10.1016/j.cell.2017.09.047;

Gompers et al. 2017: https://doi.org/10.1038/nn.4592;

Katayama et al. 2016: https://dx.doi.org/10.1038/nature19357;

Tilot et al. 2016: https://doi.org/10.1038/mp.2015.17;

Sessa et al. 2019: https://doi.org/10.1016/j.neuron.2019.07.013;

Gandal et al. 2018: https://doi.org/10.1126/science.aat8127;

Voineagu et al. 2011: https://doi.org/10.1038/nature10110;

Abrahams et al. 2013: https://doi.org/10.1186/2040-2392-4-36;

Parikshak et al. 2013: https://dx.doi.org/10.1038/nature20612;

Raw and aggregate bisulfite-seq, raw and gene-count data for RNA-seq, and raw and peak call data for ChIP seq will be available on GEO.

## Supplemental Information

Table S1. DNMT3A Mutation Table, Related to Figure 1, Figure 2, Figure S1

Table S2. BSsmooth-defined Differentially Methylated Regions, Related to Figure 4, Figure 5, Figure S5

Table S3. Table of differentially expressed genes, Related to Figure 6, Figure 7, Figure S6

Table S4. Behavioral Test Table, Related to Figure 3, Figure S3, Figure S4

